# Encoding and control of airflow orientation by a set of *Drosophila* fan-shaped body neurons

**DOI:** 10.1101/2020.07.12.199729

**Authors:** Timothy A. Currier, Andrew M. M. Matheson, Katherine I. Nagel

## Abstract

How brain circuits convert sensory signals into goal-oriented movement is a central question in neuroscience. In insects, a region known as the Central Complex (CX) is believed to support navigation, but how its compartments process diverse sensory cues to guide navigation is not fully clear. To address this question, we recorded from genetically-identified CX cell types in *Drosophila* and presented directional visual, olfactory, and airflow cues known to elicit orienting behavior. We found that a group of columnar neurons targeting the ventral fan-shaped body (ventral P-FNs) are robustly tuned for airflow direction. Unlike compass neurons (E-PGs), ventral P-FNs do not generate a “map” of airflow direction; rather they are tuned to two directions – approximately 45° to the right or left of the midline – depending on the hemisphere of the cell body. Ventral P-FNs with both direction preferences innervate each CX column, potentially forming a basis for constructing representations of airflow in various directions. We explored two possible sources for ventral P-FN airflow tuning, and found that they mostly likely inherit these responses via a pathway from the lateral accessory lobe (LAL) to the noduli (NO). Silencing ventral P-FNs prevented flies from adopting stable orientations relative to airflow in closed-loop flight. Specifically, silenced flies selected improper corrective turns following changes in airflow direction, but not after airflow pauses, suggesting a specific deficit in sensory-motor action selection. Our results identify a group of central complex neurons that robustly encode airflow direction and are required for proper orientation to this stimulus.

## Introduction

Foraging for food, locating mates, and avoiding predation all depend on an animal’s ability to navigate through complex multi-sensory environments. Many animals are known to compare and combine visual, mechanosensory and olfactory cues to achieve their navigational goals (Gire et al., 2016; Holland et al., 2009; Bianco and Engert 2015; Dacke et al., 2019; Cardé and Willis, 2008; Lockery, 2011). Identifying the brain regions and circuit organizations that support navigation with respect to different modalities is a fundamental question in neuroscience.

In insects, a conserved brain region known as the Central Complex (CX) is thought to control many aspects of navigation (Strauss and Heisenberg, 1993; Honkanen et al. 2019). The CX is a highly organized neuropil consisting of four primary subregions: the Protocerebral Bridge (PB), the Ellipsoid Body (EB), the Fan-shaped Body (FB), and the paired Noduli (NO). Columnar neurons recurrently connect these regions to each other, while tangential neurons targeting different layers of the EB and FB provide a large number of inputs from the rest of the brain (Hanesch et al., 1989; Wolff et al., 2015, Franconville et al., 2018). Outputs are primarily provided by a different subset of columnar neurons (Stone et al. 2017, Franconville et al. 2018, Xu et al. 2020).

Recent work has led to a burgeoning understanding of how the CX is functionally organized. In the EB, a group of “compass neurons” (or E-PGs) exhibit an abstract map of heading angle that is derived from both visual and airflow landmark cues (Seelig and Jayaraman, 2015; Green et al., 2017; Fisher et al., 2019; Shiozaki et al., 2020; Okubo et al., 2020). Another set of EB neurons, known as P-ENs, rotate this heading representation when the fly turns in darkness (Green et al., 2017; Turner-Evans et al., 2017). Despite these robust representations of navigation-relevant variables, genetic disruption of the EB compass network has only indirect effects on navigation. Silencing E-PGs disrupts menotaxis – straight-line navigation by keeping a visual landmark at an arbitrary angle – but not other kinds of visual orienting (Giraldo et al., 2018; Green et al., 2019).

In contrast, the FB may influence ongoing locomotor activity more directly. For example, cockroaches alter their climbing and turning strategies when the FB is lesioned (Harley and Ritzmann, 2010), while FB stimulation evokes stereotypic walking maneuvers (Martin et al., 2015). However, “compass”-like signals encoding heading and steering are also present in some parts of the FB (Shiozaki et al., 2020). Columnar neurons of the FB have been proposed to represent a desired heading, while output neurons of the FB have been proposed to drive steering (Stone et al. 2017, Honkanen et al. 2019), but these hypotheses have not been directly tested experimentally. How the FB participates in navigation, and whether its role is distinct from that of the EB, is currently unclear.

As in the EB, FB neurons represents a wide array of sensory cues, including optic flow, polarized light, and mechanical activation of the antennae or halteres (Weir and Dickinson, 2015; Heinze et al. 2009, Ritzmann et al., 2008; Phillips-Portillo, 2012; Kathman and Fox, 2019). Although it has received less attention than vision or olfaction, flow of the air or water is a critical mechanosensory cue for animals navigating in aquatic, terrestrial, and air-borne environments (Montgomery et al., 1997; Yu et al., 2016a; Alerstam et al., 2011; Reynolds et al., 2010). The primary sensors that detect flow are well-described in many species (Suli et al., 2007; Yu et al., 2016b; Yorozu et al., 2009), but an understanding of the higher brain circuits that process flow signals is just beginning to emerge (Okubo et al., 2020; Suver et al. 2019). The neurons and computations that directly support flow-based navigation remain unknown.

Here we used whole-cell recordings to systematically investigate the sensory responses of many of the major columnar cell types in the CX. We measured responses to three stimuli known to elicit basic orienting responses in *Drosophila*: a visual stripe, directional airflow, and an attractive odor. We found that columnar neurons targeting the ventral layers of the FB and the third compartment of the nodulus (“ventral P-FNs”) were robustly tuned for the direction of airflow, but not our other stimuli. Recordings from different columns suggest that ventral P-FN sensory responses are not organized in a “compass” – where all possible stimulus directions are represented as a map. Instead, ventral P-FNs primarily encode airflow arriving from two directions, approximately 45° to the right and left of the midline. Single neuron tuning depended on the hemisphere in which its cell body was located, with each column innervated by both left- and right-preferring neurons. Imaging and recording experiments suggest that this airflow representation may be inherited from the lateral accessory lobe (LAL), which projects to the third nodulus compartment (NO_3_) in each hemisphere. This anatomy could explain why all ventral P-FNs in one hemisphere share the same sensory tuning.

Genetic silencing experiments suggest that ventral P-FNs are required for normal orientation to airflow in a closed-loop flight simulator. Flies with silenced ventral P-FNs fail to make appropriate corrective turns in response to a change in airflow direction, but respond normally to airflow pauses, arguing for a specific role in linking directional sensory input to corrective motor actions. Our results support the hypothesis that different CX compartments represent sensory information in distinct formats, and identify a neural locus in the ventral FB that promotes orientation to airflow.

## Results

### Airflow dominates responses to directional sensory cues in a set of CX columnar neurons

To assess how CX compartments might differentially process sensory cues to guide navigation, we first surveyed columnar cell types that target the PB and different layers of the EB and FB. Our survey included: (1) all known columnar cell types that link the PB and NO (“P-XN” neurons); and (2) two additional cell types that target regions outside the CX proper, instead of the NO. Many of these cell types have not been previously recorded using electrophysiology. We used publicly available split-GAL4 lines (Wolff and Rubin, 2018) to express GFP in each population, then made whole-cell recordings while we presented flies with the following sensory cues, either alone or in combination: a high contrast vertical stripe, airflow generated by a pair of tubes, and apple cider vinegar, which could be injected into the airstream (Fig. 1A & 1B, right). Flies fixate vertical stripes while walking and in flight (Reichardt and Poggio, 1976; Heisenberg and Wolf, 1979; Maimon et al. 2008) and tend to orient away from an airflow source (Currier and Nagel 2018, Kaushik et al., 2020). The addition of an attractive odorant to airflow switches orientation from downwind to upwind (van Bruegel et al. 2014, Alvarez-Salvado 2018). We presented each stimulus combination from four directions: frontal, rear, ipsilateral, and contralateral (Fig. 1B, left). We monitored fly activity with an infrared camera and discarded the few trials that contained flight behavior.

**Figure 1.**
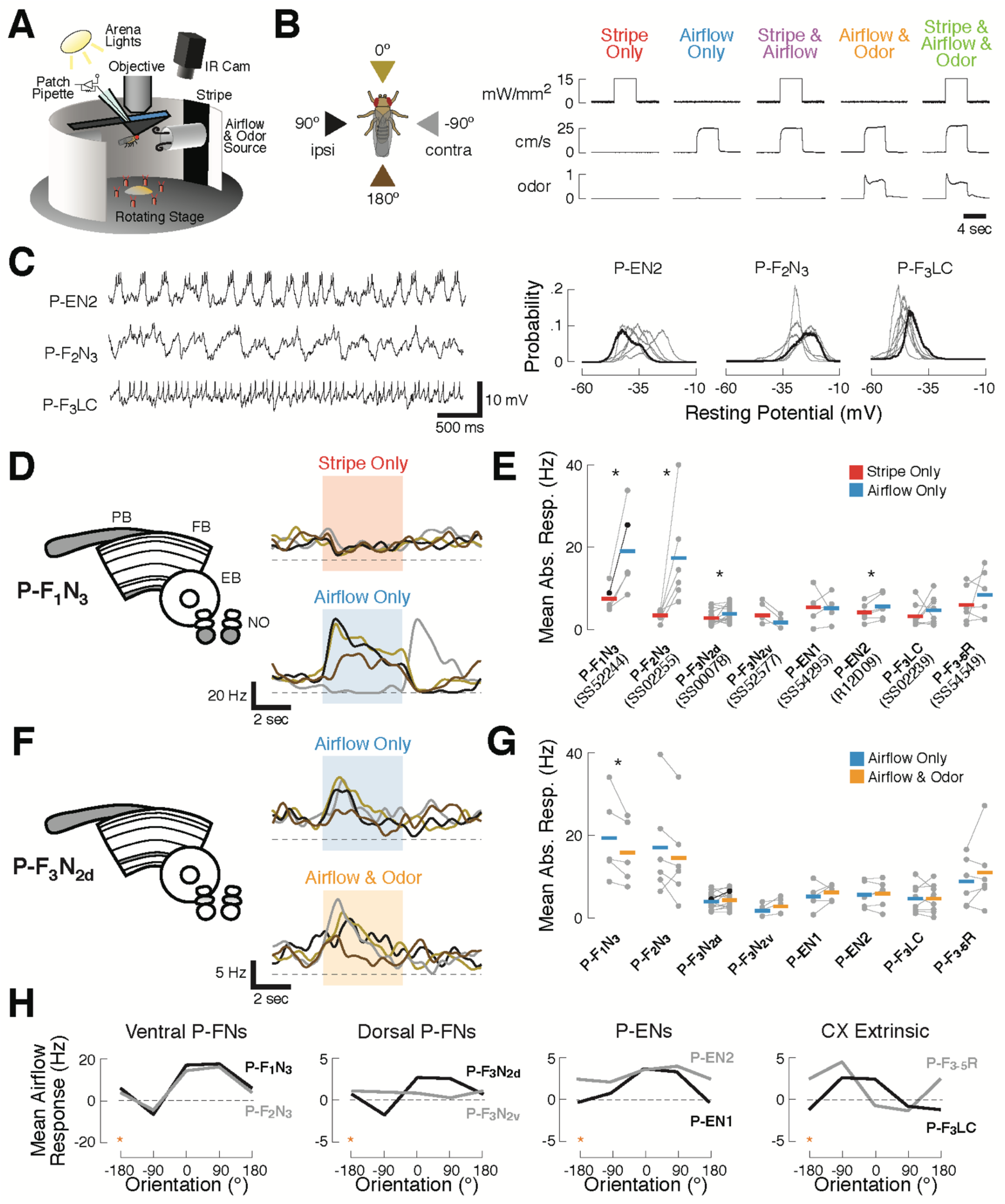
Sensory responses and preferred airflow direction vary across CX columnar cell types. (A) Experimental preparation. We targeted single neurons for patching using cell type-specific expression of GFP. Flies were placed in an arena equipped with rotatable stimulus delivery and live imaging of behavior. All data shown are from awake non-flying animals. (B) Stimulus details. Left: cue presentation directions. Front (0°, gold), rear (180°, brown), ipsilateral (90°, black), and contralateral (−90°, grey) to the recorded neuron. Right: stimulus validation. Each plot shows measurements from a photodiode (top), anemometer (middle), and photo-ionization detector (PID, bottom). PID units are arbitrary. The five stimulus combinations were: a high contrast stripe illuminated by 15 mW/cm2 ambient lighting (red), a 25 cm/s airflow stream (blue), stripe and airflow together (purple), airflow and 20% apple cider vinegar together (orange), and all three modalities simultaneously (green). Each trace is 12 sec long. Simultaneous cues were presented from the same direction. (C) Rhythmic and tonic baseline activity in a subset of CX columnar neuron types. Left: raw membrane potential over time for three example neurons. P-EN2 and P-F2N3 show rhythmic activity at different frequencies, while P-F3LC fires tonically at rest. Right: resting membrane potential probability distributions for each recorded neuron of the types shown (gray). Example neurons in black. Rhythmic neurons exhibit broad distributions, while tonic neurons show tight distributions. See also Fig. S1. (D) Left: CX neuropils innervated by P-F1N3 neurons (gray). PB, Protocerebral Bridge; FB, Fan-shaped Body; EB, Ellipsoid Body; NO, noduli. Right: PSTHs for a single P-F1N3 neuron. Each trace represents the mean of four presentations of stripe alone (red, top) or airflow alone (blue, bottom) from one direction. Colors representing different directions as illustrated in (B). Colored boxes indicate the 4 sec stimulus period. Dashed line indicates 0 Hz. (E) Responses to airflow (blue) versus stripe (red) for each neuron type. Gray dots indicate the mean spiking response of each cell (1 sec stimulus minus 1 sec baseline) to four trials from the direction producing the strongest response (see Methods). Colored bars: mean across cells. The example P-F1N3 neuron from (D) is shown in black. Significant differences (*p* < 0.05 by rank-sign test) between modalities are marked with an asterisk. For additional detail, see Fig. S2. (F) Left: CX neuropils innervated by P-F3N2d neurons. Right: PSTHs for a single P-F3N2d neuron. Each trace represents the mean of four presentations of airflow alone (blue, top) or airflow and odor together (orange, bottom) from one direction. Plot details as in (D). (G) Responses to odorized airflow (orange) versus airflow alone (blue) for all cell types recorded. Asterisk: odor significantly reduces the response of P-F1N3. Plot details as in (E). For additional detail, see Fig. S2. (H) Mean airflow response across cells as a function of airflow direction for each cell type. Cell types are plotted in groups of two (gray, black) according to broad anatomical similarities. Note different vertical axis scales. Data at −180° is replotted from 180° for clarity (orange stars). For additional detail, see Fig. S2.

We first noticed that CX columnar cell types possessed diverse baseline activity (Fig. S1 and Table 1). Some cell types, such as P-EN2 and P-F2N3, showed rhythmic fluctuations in membrane potential, evident in the timecourses and distributions of membrane potential (Fig. 1C). Rhythmic neurons exhibited broader membrane potential distributions than less rhythmic neurons (Fig. 1C, right and Fig. S1B&C). P-F3LC, for example, showed stable baseline activity and a lower characteristic resting potential. We did not observe any correlation between fly behavior and the presence or absence of oscillations, but resting membrane potential did rarely fluctuate with leg movements. Input resistance varied by cell type, with values ranging from 1.5 - 10 GOhm (Fig. S1D).

**Table 1.** Intrinsic properties of surveyed neuron types. Resting potential, input resistance, and characteristic oscillatory frequency are shown for each recorded cell type. Values represent the cross-fly mean +/- SEM. See also Fig. S1.

| Cell Type | Driver Line | N | Resting Potential (mV) | Input Resistance (GΩ) | Osc. Freq. (Hz) |
| --- | --- | --- | --- | --- | --- |
| P-F1N3 | SS52244 | 6 | -18.0 ± 1.0 | 6.21 ± 0.52 | 4 |
| P-F2N3 | SS02255 | 6 | -22.0 ± 1.3 | 4.86 ± 0.50 | 2 |
| P-F3N2d | SS00078 | 14 | -30.9 ± 0.9 | 2.58 ± 0.24 | - |
| P-F3N2v | SS52577 | 4 | -31.8 ± 3.5 | 6.00 ± 0.50 | - |
| P-EN1 | SS54295 | 4 | -32.9 ± 2.1 | 2.52 ± 0.29 | 3 |
| P-EN2 | R12D09 | 6 | -30.1 ± 1.9 | 3.30 ± 0.29 | 3 |
| P-F3LC | SS02239 | 8 | -39.7 ± 0.8 | 2.75 ± 1.11 | - |
| P-F3-5R | SS54549 | 6 | -26.3 ± 1.3 | 7.89 ± 0.51 | - |
| E-PG | SS00090 | 4 | -29.4 ± 1.5 | 2.30 ± 0.35 | - |

We next turned our attention to the sensory responses of CX columnar cell types. We found that, although most neurons responded to each cue in some manner, airflow responses were generally larger than stripe responses across cell types in the survey. In P-F_1_N_3_ neurons, for example, large directionally tuned spiking responses were observed during airflow presentation, but not stripe presentation (Fig. 1D). To assess this difference across cell types, we identified the cue direction(s) that elicited the largest stripe and airflow responses for each recorded neuron. We then took the mean absolute value of responses to that direction (relative to baseline), and plotted these values as a function of sensory condition for each cell (Fig. 1E). Airflow responses were largest in two cell types targeting the ventral layers of the FB and the third compartment of the NO: P-F_1_N_3_ and P-F_2_N_3_. However, the cross-fly mean airflow response was larger than the mean stripe response in all cell types except P-F_3_N_2v_ and P-EN1. This difference was significant for half of the cell types surveyed: P-F_1_N_3_, P-F_2_N_3_, P-F_3_N_2d_, and P-EN2. The strength of sensory tuning tended to vary with raw response magnitude, with robustly responsive cell types also showing the strongest directional preferences (Fig. S2). As such, many of the neurons we recorded were tuned for the direction of airflow (Fig. 1H).

In contrast to this strong directional preference in the airflow condition, tuning strength in the visual condition was relatively weak across recorded cell types (Fig. S2). One notable exception were P-ENs, which displayed modest visual tuning, in agreement with previous results (Green et al., 2017; Fisher et al., 2019). To ensure that our visual stimulus was sufficient to evoke responses in the CX, we recorded a small number of E-PG neurons, which are known to be tuned for both stripe and airflow direction (Seelig and Jayaraman, 2015; Green et al., 2017; Okubo et al., 2020). Recorded E-PGs showed directional tuning to the stripe (Fig. S3), indicating that our stimulus can evoke tuned visual responses.

Adding odor to the airflow stream had only mild effects on CX columnar neuron responses. Odor slightly reduced the airflow responses of P-F_1_N_3_ and P-F_2_N_3_ (Fig. 1G), although this difference was only significant for P-F_1_N_3_. Some single cells also showed increases or decreases in spiking activity in the odor condition (Fig. 1F). While response magnitudes no doubt depend on stimulus intensity (airflow velocity, stripe contrast, and odor concentration), we know that our visual and airflow cues elicit orienting responses of approximately equal magnitude in a flight simulator (Currier and Nagel, 2018). Similarly, our odor cue produces robust orientation changes in walking flies (Alvarez-Salvado et al. 2018). Therefore, these responses reflect differential neural encoding of stimuli with similar behavioral relevance.

Broadly, our survey suggested that pairs of columnar cells innervating the same NO compartment share similar sensory responses. In particular, “ventral P-FNs,” which receive input in layer 3 of the NO and ventral FB layers, had large airflow responses, showed olfactory suppression, and preferred ipsilateral airflow (Fig. 1H). “Dorsal P-FNs,” which receive input in layer 2 of the NO, and “P-ENs,” which receive input in layer 1 of the NO, both displayed modest sensory responses and ipsilateral airflow tuning in most cases. In contrast, the “CX extrinsic” columnar neurons we recorded, which target neuropils outside the CX, both preferred contralateral airflow. Thus, our survey suggests that sensory responses in CX neurons vary according to their input neuropils.

### Multi-sensory cues are summed in CX neurons, with some layer-specific integration variability

If CX compartments process unique combinations of sensory signals, we reasoned that neurons targeting different layers might also integrate multi-sensory signals in distinct ways. To understand the computational principles that govern cue integration in our recordings, we first compared multi-sensory responses to the sum of single modality responses (Fig. 2A). Summation is a simple circuit principle that is naturally achieved by an upstream neuron or neurons passing multimodal cues to a single downstream cell. For each cell type, we found the mean response to the airflow-plus-stripe condition for each cue direction. We then plotted these multi-sensory responses against the sum of the mean airflow response and the mean stripe response from the same directions (Fig. 2A, left). When we plotted these measures for each cell type, we found that most points fell on or near the diagonal, indicating that multi-sensory responses are, on average, approximately equal to the sum of single modality responses. This trend of near-perfect summation was also true when all three modalities were presented simultaneously (Fig. 2A, right). These results suggest that poly-modal integration generally proceeds via a summation principle, at least for the stimuli presented here, which were always presented from the same direction.

**Figure 2.**
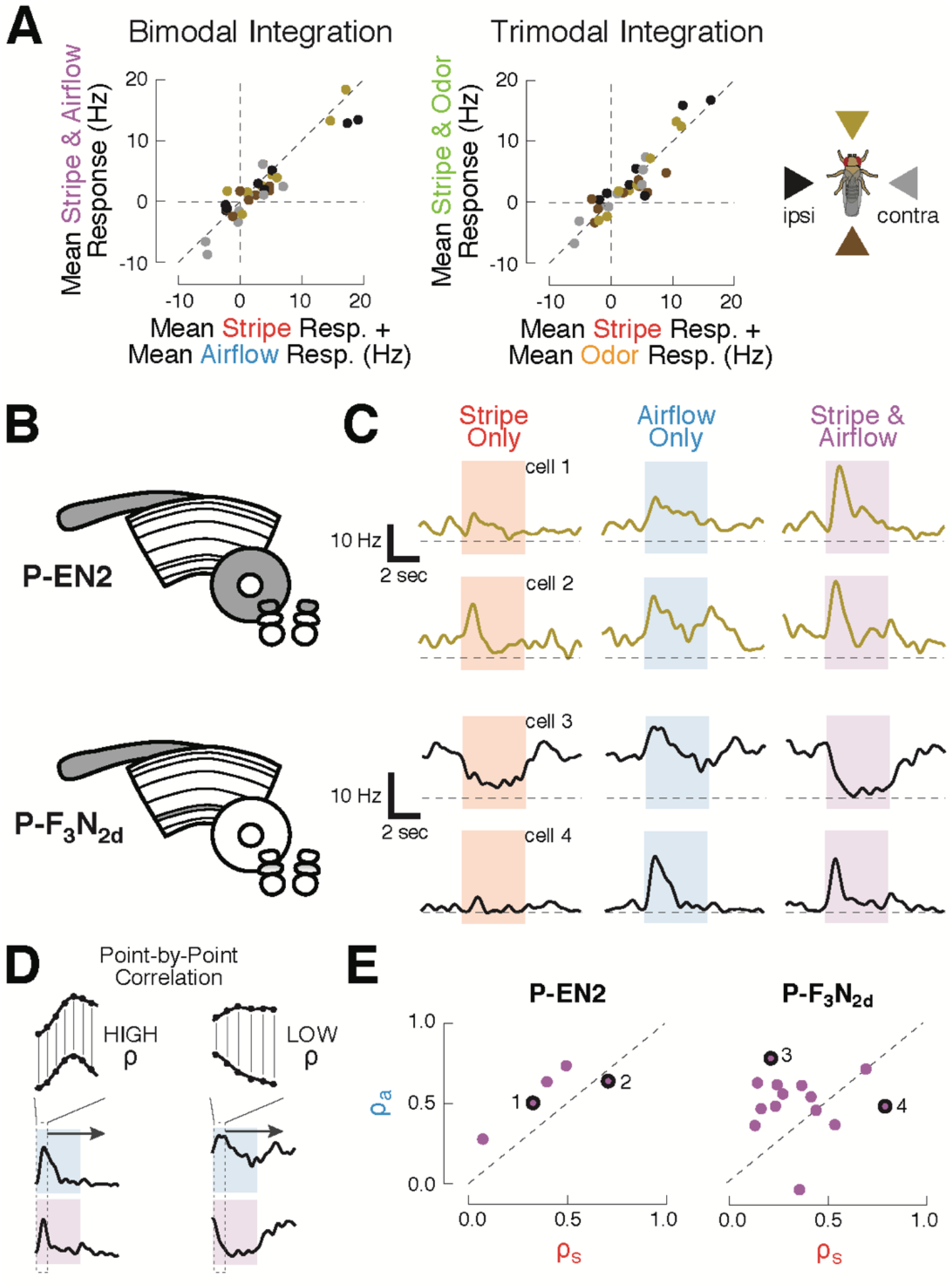
CX columnar neurons sum inputs from different modalities on average, but show diverse integration strategies at the level of single cells. (A) Summation of multimodal cues. Left: mean spiking response to stripe & airflow together versus sum of mean stripe alone and airflow alone responses. Each point represents the response of one cell type to cues from one direction. Right: mean spiking response to stripe & airflow & odor versus sum of mean stripe alone response and mean airflow & odor response. Colors indicate cue direction (far right). Data falling along the diagonals indicate perfectly weighted summation. (B) CX neuropils innervated by example cell types P-EN2 and P-F3N2d. (C) PSTHs of two neurons from each cell type. Curves represent mean firing rate across four trials of each stimulus from a single direction. Colored boxes indicate the four second stimulus period. Dashed lines indicate 0 Hz. Top: example P-EN2 neurons responding to frontal cues. In both cases, the multi-sensory response (purple) is a weighted sum of the single modality responses (red, blue). Bottom: P-F3N2d neurons responding to ipsilateral cues. In one cell (cell 3) the stripe response dominates the multi-sensory response, while in the other (cell 4) the airflow response dominates. (D) Correlation method for computing response similarity. We computed a point-by-point correlation between the mean baseline-subtracted firing rate timecourses of multi-sensory (airflow and stripe together) responses and responses to a single modality (ariflow alone, or stripe alone), across all stimulus directions. Similar traces result in high correlation coefficient (ρ). (E) Correlation coefficients (calculated as in D) of the multimodal response (stripe and airflow together) to each single modality response (airflow alone, ρa, or stripe alone, ρs). Data along the diagonal indicates that the multi-sensory response is equally similar to the stripe alone and airflow alone responses, a hallmark of summation. Data off diagonal indicates that one modality dominates the multi-sensory response. The four example cells from (C) are labeled with numbers and black rings. P-EN2 neurons consistently sum stripe and airflow responses (top), while P-F3N2d neurons integrate with greater diversity (bottom).

Because of the diverse sensory responses observed in our survey, we wondered whether integration principles may also differ across individual members of a single cell type. To answer this question, we evaluated stripe and airflow integration for single cells (Fig. 2B-E). Broadly, we found that integration diversity was small for some cell types, but large for others. P-EN2 neurons, for example, showed remarkably consistent summation. Single neuron spiking responses to multi-sensory cues strongly resembled the responses to single modality cues across all P-EN2s (Fig. 2C, top). To evaluate summation in a scale-free manner, we found the correlation coefficient between the mean response timecourse to the multi-sensory condition (airflow and stripe together) and each of the single modality conditions (airflow alone or stripe alone, Fig. 2D). To do this, we concatenated each cell’s mean spiking responses to stimuli presented from different directions (−90°, 0°, 90°, 180°), with the baseline period removed. We then took the point-by-point correlation between the concatenated multi-sensory response and the single modality responses, which yielded a pair of correlation coefficients (ρ_a_ for airflow and ρ_s_ for stripe). A coefficient of 1 indicates that the multi-sensory and single modality data varied over time in perfect synchrony. Conversely, a coefficient closer to 0 indicates that these signals did not vary together. When we plotted these coefficients against one another (Fig. 2E, but also see Fig. S4), we found that the P-EN2 data lie along the diagonal, indicating that the stripe and airflow responses equally resemble the multi-sensory response for each neuron.

In contrast, P-F_3_N_2d_ neurons show much greater integration diversity. While some cells displayed multi-sensory activity that was dominated by the stripe response, others were dominated by the airflow response (Fig. 2C, bottom). Indeed, the correlation coefficients for P-F_3_N_2d_ reveal a full range of modality preferences (Fig. 2E, right). We did not observe any obvious relationship between single neuron anatomy and the method of integration used by that cell, although this result might reflect a limitation of our stimuli — for example, if these neurons were preferentially tuned to stimuli at a particular phase offset. Thus, while summation appears to govern P-F_3_N_2d_ integration on average, individual neurons show an array of sensory integration strategies. This trend of summation on average, but diversity at the single cell level, was found for many of the surveyed cell types (Fig. S4). These results suggest that CX neurons integrate multi-modal sensory cues with compartment-specific variability.

### Ventral P-FN airflow responses are organized as orthogonal basis vectors, rather than as a map or compass

At the conclusion of our survey, ventral P-FNs (P-F_1_N_3_ and P-F_2_N_3_) stood out as possessing the most robust sensory responses, prompting us to examine the activity of P-F_2_N_3_ in greater detail (Fig. 3A-G). These cells had resting membrane potentials between −30 and −25 mV, and input resistances around 3 GOhm (Fig. S1). P-F_2_N_3_s showed strong spiking responses to single presentations of ipsilateral airflow, and active inhibition followed by offset spiking during single presentations of contralateral airflow (Fig. 3B&D). Spiking activity in response to ipsilateral and contralateral airflow was relatively consistent from trial to trial, while frontal and rear airflow elicited more diverse responses relative to baseline on each trial (Fig. 3D). On average, P-F_2_N_3_ neurons showed graded membrane potential and spiking responses to airflow, but not to the stripe (Fig. 3C&E).

**Figure 3.**
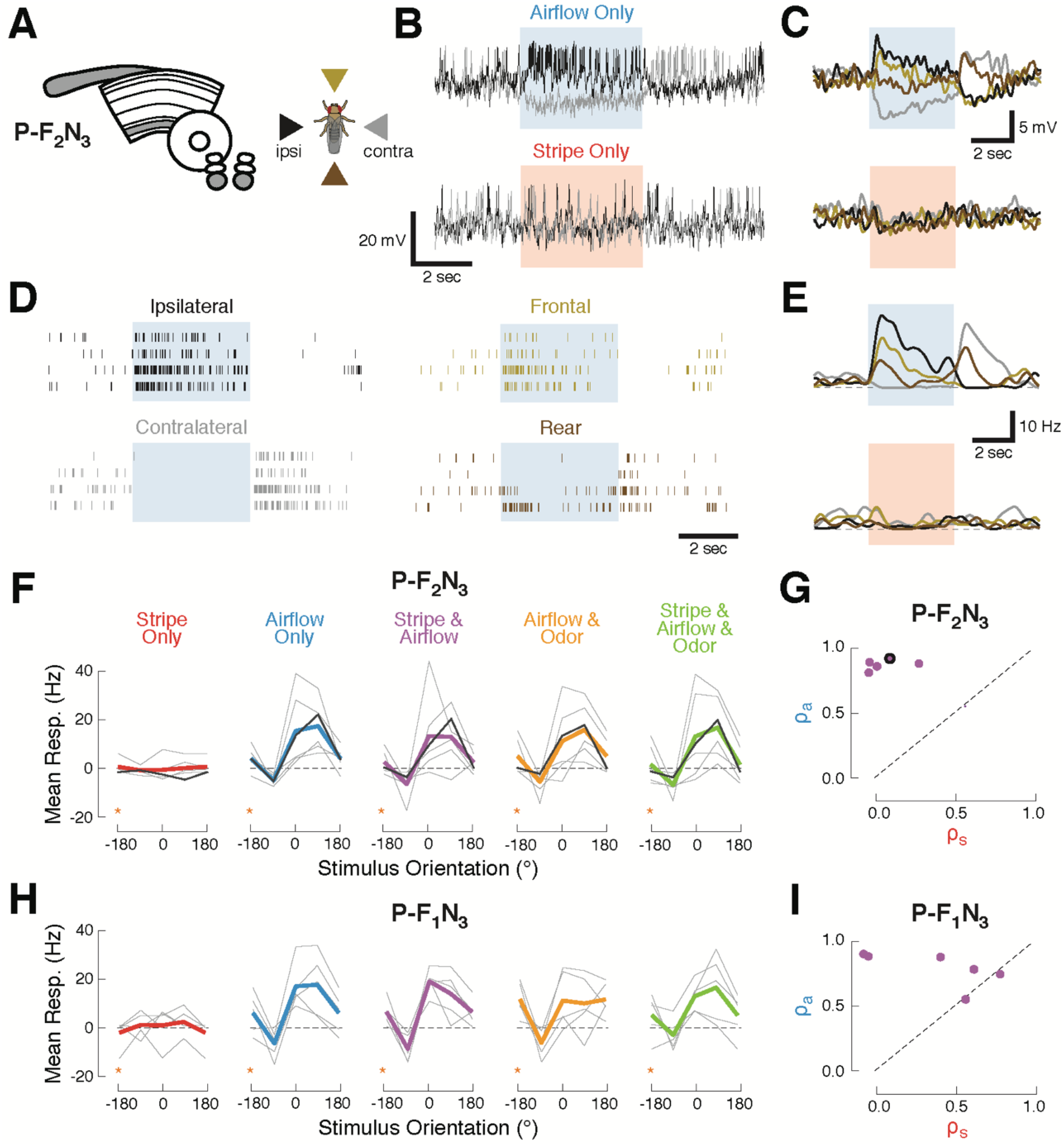
Ventral P-FNs selectively respond to directional airflow. (A) Left: CX neuropils innervated by P-F2N3. Right: color key for directional stimuli. (B) Example trials from a single P-F2N3 neuron. Raw membrane potential for single presentations of airflow alone (blue, top) or stripe alone (red, bottom) for ipsilateral (black) and contralateral (gray) directions. Colored box indicates 4 sec stimulus period. Baseline Vm = −28 mV. (C) Average Vm (over 4 trials) for the example neuron shown in (B). Colors represent directions as shown in (A). Stimulus period represented as in (B). (D) Spike response rasters for the example neuron shown in (B). Colors and stimulus period as in (C). (E) PSTHs for the example neuron in (B). Colors and stimulus period as in (C). (F) P-F2N3 direction tuning for each cue set showing that responses to airflow are not modulated by other modalities. Mean spiking response minus baseline for each recorded cell as a function of stimulus direction (gray lines). The example neuron in (B-E) is shown in black. Mean tuning across cells shown in thick colored lines. Data at −180° is replotted from 180° for clarity (orange stars). (G) Similarity (as in Fig. 2C) of P-F2N3 multi-sensory (stripe + airflow) responses to airflow alone and and stripe alone. Response to stripe + airflow is highly similar to airflow alone. Example neuron marked in black. (H) Direction tuning for the second type of ventral P-FN, P-F1N3. Note that odor subtly inhibits airflow-evoked responses (as shown in Fig. 1G). Data at −180° is replotted from 180° for clarity (orange stars). (I) Same as (G), but for P-F1N3.

To assess how additional sensory modalities modulate this directional airflow tuning, we plotted the spiking response as a function of stimulus orientation for each combination of cues (Fig. 3F). We found that mean tuning across the population did not change when the stripe, odor, or both, were added to airflow. Each P-F_2_N_3_ neuron showed a large airflow correlation coefficient and a small visual coefficient (Fig. 3G), indicative of multi-sensory responses that strongly resemble the airflow-only response. P-F_1_N_3_ sensory activity was similar, except for the olfactory suppression noted above (Fig. 3H&I).

Like many other columnar cell types, individual ventral P-FNs target one CX column, with cell bodies in each hemisphere collectively innervating all eight columns (Wolff et al., 2015). We next asked whether neurons innervating different columns show distinct directional tuning, as has been previously observed for polarized light cues (Heinze and Homberg, 2007) and for visual landmarks and airflow in E-PG compass neurons (Green et al., 2017; Fisher et al. 2019, Okubo et al. 2020). To address this question, we recorded from a larger set of P-F_2_N_3_ neurons while explicitly attempting to sample from a range of CX columns (Fig. 4A). For this experiment, we presented airflow from eight directions and omitted other sensory modalities. To identify the FB columns targeted by our recorded neurons, we filled each cell with biocytin and visualized its anatomy after the recording session (Fig. 4B&C).

**Figure 4.**
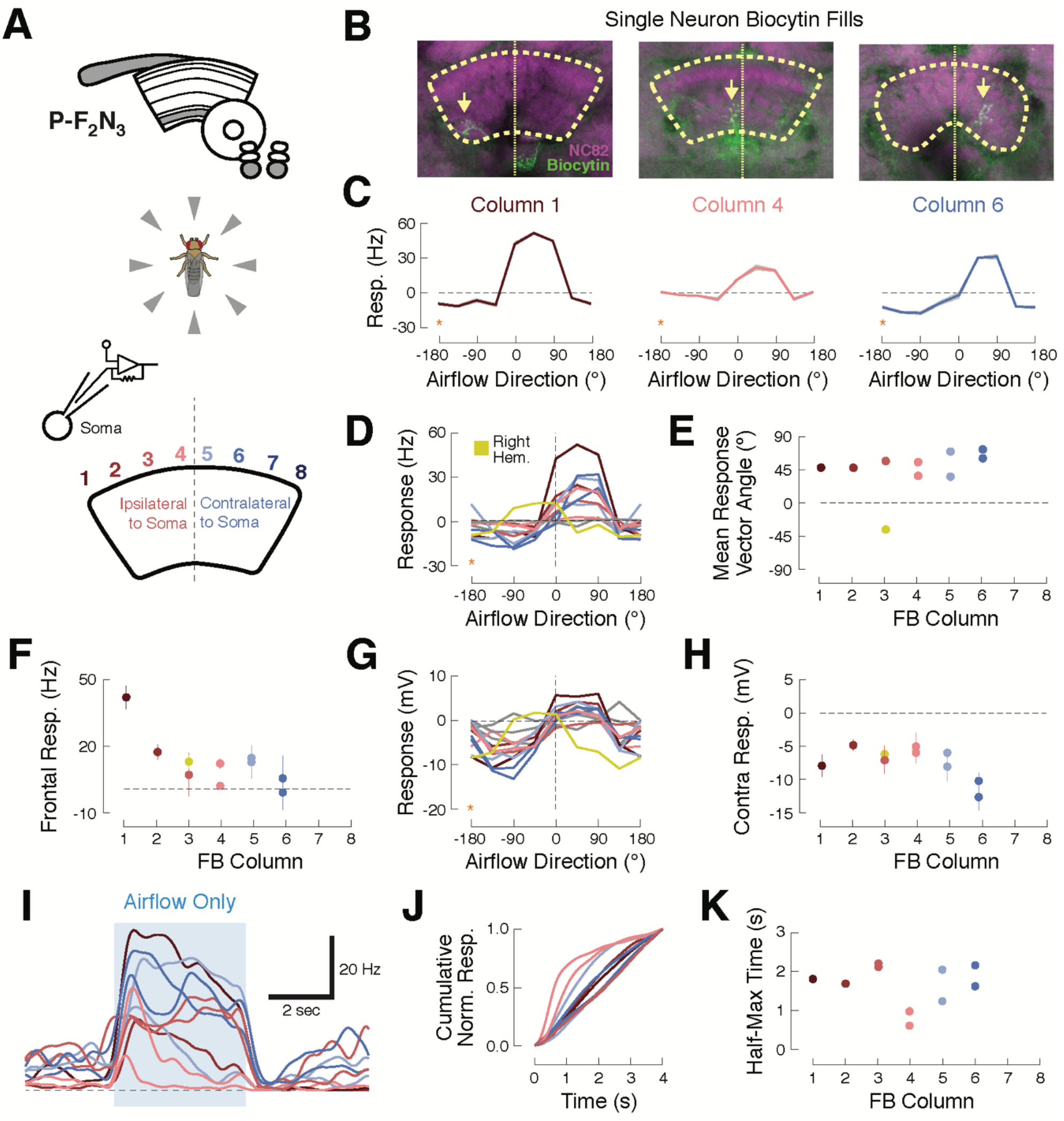
Ventral P-FNs exhibit similar ipsilateral airflow tuning across CX columns. (A) Top: CX neuropils innervated by P-F2N3. Bottom: experimental setup. We presented airflow from eight directions and identified the column innervated by each patched neuron by filling the cell with biocytin. (B) Biocytin fills (green) for three example cells innervating columns 1 (left), 4 (middle), and 6 (right). Yellow arrows indicate FB portions of fills. Neuropil in magenta. Thick dashed line indicates the borders of the FB and thin dotted line shows the midline. (C) P-F2N3 airflow tuning is similar across FB columns. Mean +/- SEM spiking response as a function of airflow direction for the three example cells shown in (B). Data at −180° is replotted from 180° for clarity (orange stars). Colors reflect innervated column, as in (A). (D) Mean spiking response as a function of airflow direction for all recorded P-F2N3 neurons. Colors as in (A). A single right hemisphere neuron is shown in yellow. Gray curves indicate cells for which no anatomy data could be recovered. Data at −180° is replotted from 180° for clarity (orange star). (E) Mean response vector angle as a function of column for each recorded neuron. Colors as in (D). (F) Mean +/- SEM spiking response to frontal airflow as a function of column. Colors as in (A). (G) Mean membrane potential response as a function of airflow direction for each neuron. Data at −180° is replotted from 180° for clarity (orange star). Colors as in (D). (H) Mean +/- SEM membrane potential response to contralateral airflow as a function of column. Colors as in (D). (I) Timecourse (PSTH) of airflow responses to ipsilateral airflow for the same P-F2N3 neurons (average of 4 trials). Blue box indicates 4 sec stimulus. Colors as in (A). (J) Cumulative normalized response for each neuron during the 4 sec stimulus, normalized to its mean integrated response. Transient responses show fast rise times and tonic responses show slower rise times. Colors as in (A). (K) Time to half-max (a measure of response transience) as a function of column. Colors as in (A).

Surprisingly, we found that all left-hemisphere P-F_2_N_3_ neurons responded strongly to airflow presented from the front-left (between 0° and 90°, ipsilateral) and were inhibited by airflow from the rear-right (between −90° and −180, contralateral), regardless of the column they innervated. One right hemisphere neuron was excited by airflow from the front-right and inhibited by airflow from the rearleft. Thus, all P-F_2_N_3_ neurons show a preference for airflow presented ipsi-frontal relative to the hemisphere of their cell bodies (Fig. 4D&E), with peak tuning around approximately 45° ipsilateral.

We did notice some subtle differences in tuning that varied with column. The spiking response to frontal airflow (0°) was strongest in the ipsilateral-most column 1, and was weaker in more contralateral columns (Fig. 4F). Membrane potential responses to contralateral also airflow varied by column, with contralateral columns exhibiting the greatest inhibition relative to baseline (Fig. 4G,H). P-F_2_N_3_ neurons also showed temporally diverse airflow responses (Fig. 4I). Some neurons showed sustained activity during the stimulus period, while others displayed only transient responses to airflow presentation. Temporal responses were not reliably organized by column (Fig. 4J,K) but might instead reflect different behavioral states of the animal.

These data support two conclusions. First, ventral P-FNs respond primarily to airflow, and not to the other stimuli presented in our cue set. Second, airflow tuning across P-F_2_N_3_ neurons is not organized as a map of airflow direction, but is instead clustered around two directions approximately 45° to the left and right of the fly midline. This is reminiscent of the organization of optic flow responses in TN neurons of the bee (Stone et al. 2017), which have been proposed to act as basis vectors for computing movement of the animal through space. Since each FB column is innervated by ventral P-FNs with cell bodies in both hemispheres, each column receives two orthogonal airflow signals that could be used to construct tuning to a variety of airflow directions in downstream neurons.

### Ventral P-FNs likely inherit their airflow tuning from the Lateral Accessory Lobe (LAL)

What is the source of the airflow signals in ventral P-FNs? Neurons sensitive to airflow direction have recently been identified in both the antler (ATL, Suver et al., 2019), and the lateral accessory lobe (LAL, Okubo et al., 2020). Using trans-tango experiments (not shown), and the *Drosophila* hemibrain connectome (Xu et al., 2020), we identified neurons presynaptic to ventral P-FNs that receive input in each of these regions (Fig. 5A). Ventral FB neurons (vFBNs) receive input in the antler and project to layer 2 of the FB, while LNa neurons (LAL-NO(a) neurons, Wolff and Rubin, 2018) receive input in the LAL and project to the third compartment of the NO. To assess whether either of these groups of neurons carry tuned airflow signals, we recorded from vFBNs and performed 2-photon calcium imaging using GCaMP6f from LNa neurons (Fig. 5B). LNa somata were not accessible for electrophysiology in our preparation.

**Figure 5.**
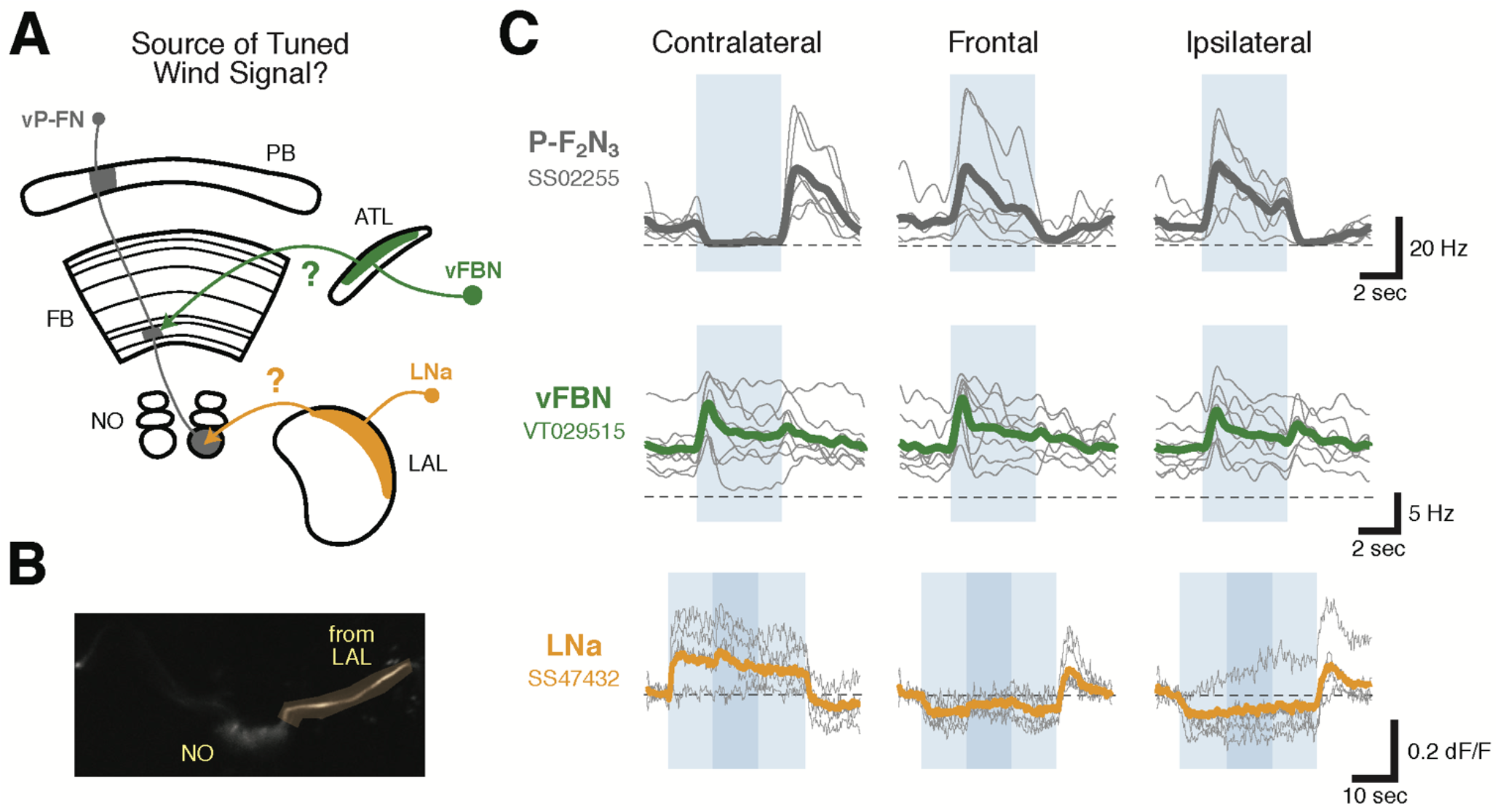
LNa neurons are a likely source of airflow signals in ventral P-FNs. (A) Experimental framework. Neurons with tuned airflow responses have recently been identified in the Antler (ATL) and Lateral Accessory Lobe (LAL). We recorded from vFBN (green) and LNa (orange) neurons to assess which input pathway might carry tuned airflow signals. (B) Single plane image of the NO region of the *SS47432 > UAS-GCaMP6f* line used to record LNa calcium activity. Imaging ROI, highlighted in orange, is the neurite of one LNa neuron in one hemisphere that connects the LAL and NO. (C) Ventral P-FN airflow tuning is likely inherited from the LAL. Mean firing rate (top two rows) or dF/F (bottom row) as a function of time is plotted for each fly (thin gray lines). Cross-fly mean activity is plotted as thick colored lines. Responses to airflow presented contralaterally (−90°, left column), frontally (0°, middle column), and ipsilaterally (90°, right column) are shown. Directions (ipsi, contra) are relative to the hemisphere of connected ventral P-FN cell bodies. vFBNs responded to airflow, but were not sensitive to airflow direction. LNa neurons showed strong directional tuning for airflow that is sign-inverted with respect to ventral P-FN activity. Blue boxes represent stimulus period (4 sec for top two rows, 30 sec for bottom row), while dashed lines indicate 0 Hz or dF/F. Darker blue region in the bottom row represents a 10 sec period when 10% apple cider vinegar was injected into the airstream (while maintaining constant airflow velocity). Odor did not have a statistically significant impact on LNa activity. Colors as in (A).

We found that LNa neurons, but not vFBNs, possessed directionally tuned airflow responses (Fig. 5C). Like ventral P-FNs, LNa neuron activity was strongly modulated by wind direction but not by the presence of odor (Fig. 5C). Since all of the strongly airflow-tuned ventral P-FNs receive input in the third compartment of the NO in one hemisphere, this finding could explain why all ventral P-FNs in one hemisphere share similar sensory tuning. Right-hemisphere LNa’s are connected to left-hemisphere ventral P-FNs, and vice-versa (Fig. 5A). When we specifically compared the tuning of LNa neurons to synaptically connected ventral P-FNs, we found that LNa tuning was inverted (Fig. 5C), suggesting that either LNa neurons are inhibitory, or ventral P-FNs do not *directly* inherit their tuning from LNa neurons. Together, these results suggest that tuned airflow responses in ventral P-FNs are likely inherited from airflow-sensitive populations in the LAL (WL-L neurons, Okubo et al., 2020), although silencing experiments will be required to directly test the contribution of LNa and WL-L neurons to ventral P-FN sensory tuning.

### Ventral P-FNs are required to orient to airflow in tethered flight

Finally, we wondered whether ventral P-FNs play a role in orientation to airflow. We addressed this question with a previously designed closed-loop flight simulator that uses an infrared camera to monitor the fictive turning of a tethered animal flying in the dark (Fig. 6A). This turn signal drives rotations of an airflow tube, allowing flies to control their orientation with respect to that flow. In previous experiments, we observed that flies prefer to orient away from the source of flow (Currier and Nagel, 2018).

**Figure 6.**
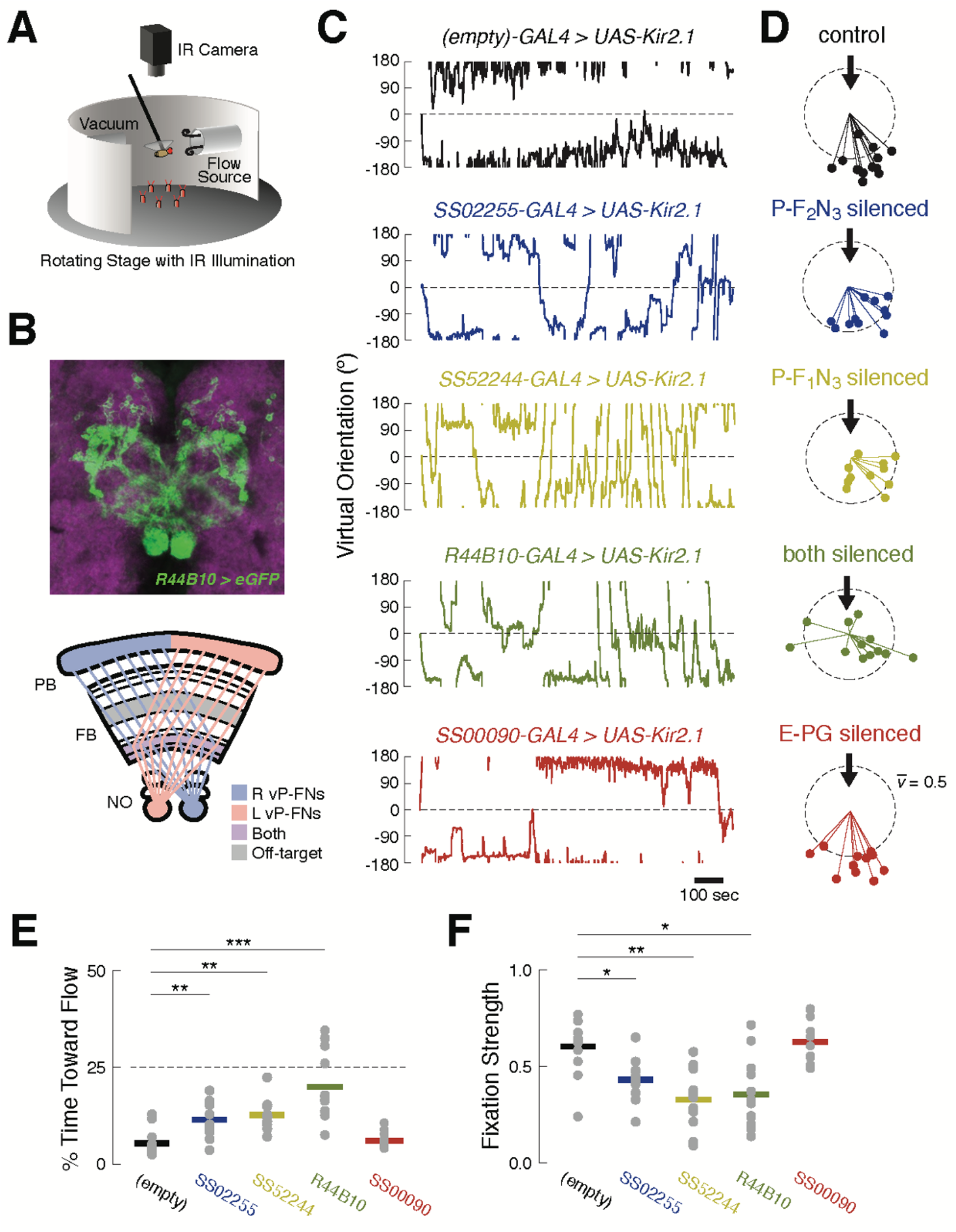
Silencing ventral P-FNs disrupts orientation to airflow. (A) Schematic of flight simulator arena. Rigidly tethered flying flies orient in closed-loop with an airflow stream. Infrared illumination is used to track wingbeat angles which drive airflow rotation. The arena was otherwise in darkness. Modified from Currier and Nagel (2018). (B) Anatomy of *R44B10-GAL4*, a driver line that targets both P-F1N3 and P-F2N3. Top: maximum z-projection of R44B10 driving *10XUAS-Kir.21-eGFP* (green). Neuropil in magenta. Bottom: schematic of CX neuropils labeled by *R44B10-GAL4*. Right hemisphere ventral P-FNs (blue) target the right half of the PB, the entire FB, and the left NO. Left hemisphere ventral P-FNs (red) target the left half of the PB, the entire FB, and the right NO. Neurons from both hemispheres target all columns of FB layers 1 and 2 (purple). *R44B10-GAL4* also labels non-P-FN neurons in a FB layer (“off target,” grey). Whole-brain expression in Fig. S5. (C) Orientation over time for example flies of each genotype. Control flies (*empty-GAL4 > UAS-Kir2.1*, black) fixate orientations away from the airflow sources. This fixation is reduced in flies with P-F2N3 (blue), P-F1N3 (yellow), or both (dark green) silenced. Flies with EPGs silenced (red) show control-like orientation behavior. The entire 20 min testing period is shown for each fly. Airflow emanated from 0° (dashed line). (D) Stick-and-ball plots of mean orientation (ball angle) and fixation strength (stick length) for each fly tested in the airflow orienting paradigm. Fixation strength is the length of the mean orientation vector, which is inversely proportional to circular variance (see Methods). Dashed circle corresponds to fixation strength of 0.5. All but one control fly, and all E-PG-silenced flies, showed fixation strengths near or above this value, while ventral P-FN-silenced flies displayed smaller fixation strengths. Thick arrow signifies the position and direction of the airflow stimulus (0°). Colors as in (C). (E) Percentage of time each fly (gray dots) oriented toward the flow source (between +45° and −45°), as a function of genotype. Horizontal bars indicate cross-fly means, with colors as in (C). Dashed line indicates the expected value for random orienting (chance). **, *p* < 0.01; ***, *p* < 0.001 (rank-sum test). (F) Fixation strength (as illustrated in (D)) for each fly as a function of genotype. A value of 1 indicates perfect fixation. Plot details as in (E). *, *p* < 0.05; **, *p* < 0.01 (rank-sum test).

We first asked whether silencing ventral P-FNs (P-F_1_N_3_ and P-F_2_N_3_) impairs normal airflow-based orienting. We compared behavior in flies where these neurons were silenced with Kir2.1 to control flies where Kir2.1 was driven by an *empty-GAL4* cassette. Consistent with our previous results, control flies adopted stable orientations away from the airflow source (Fig. 6C-F). When we calculated the mean orientation vector for each control fly, we found that they all preferred orientations roughly opposite the flow source, near 180° (Fig. 6D). When either type of ventral P-FN was silenced with Kir2.1, flies displayed partially impaired orientation ability (Fig. 6D). These groups showed increased orienting toward the flow (Fig. 6E) and reduced orientation stability (Fig. 6F) compared to controls, although both effects were moderate.

Given the similar sensory responses of P-F_1_N_3_ and P-F_2_N_3_, we reasoned that they might serve overlapping roles. We therefore sought a driver line that labeled both classes of ventral P-FN, and identified *R44B10-GAL4* as one such line with minimal off-target expression (Figs. 6B and S5). Silencing *R44B10-GAL4* neurons with Kir2.1 produced a more severe phenotype (Fig. 6C). Compared to controls, these flies showed less stable orientation to airflow (Fig 6D), spent significantly more time oriented toward the airflow source (Fig. 6E), and had reduced fixation strength (Fig. 6F). Collectively, these results suggest that ventral P-FNs are required for normal orientation to airflow. We attempted to broadly silence ventral P-FNs using several other genotypes (*15E12-GAL4, 67B06-GAL4*, and *20C08-GAL4*), however none of these flies were viable when crossed to *UAS-Kir2.1*.

E-PG neurons also respond robustly to directional airflow cues (Okubo et al. 2020). Thus, we wondered whether silencing these cells would also impair orientation to airflow. In contrast to our experiments with ventral P-FNs, we found that silencing E-PGs did not disrupt orientation to airflow (Fig. 6C-F). In flies with silenced E-PGs, both the time spent orienting toward the airflow, and orientation stability, were indistinguishable from controls (Fig. 6E-F). The split-GAL4 line used to silence E-PGs was similarly sparse to the lines used to silence P-F_1_N_3_ and P-F_2_N_3_ alone, strengthening the conclusion that ventral P-FNs specifically play a functional role in orientation to airflow.

To obtain more insight into the mechanism by which ventral P-FNs control orientation, we pseudorandomly introduced six stimulus perturbations throughout each 20 minute closed-loop testing session (Fig. 7A). These were long (2 sec) or short (150 ms) pauses in airflow (“airflow off”), and long (63.36°) or short (14.44°) angular displacements of the airflow source to the left or right (airflow “slip”). In response to airflow pauses, control flies briefly turned towards the airflow source (Fig. 7B). When the airflow resumed, flies once again turned away (see also Currier and Nagel. 2018). Overall turning in response to airflow pauses remained unchanged when both types of ventral P-FN were silenced (Fig. 7C). These results suggest that, despite the abnormal orienting behavior shown by ventral P-FN-silenced flies, they are still able to detect airflow and determine its direction.

**Figure 7.**
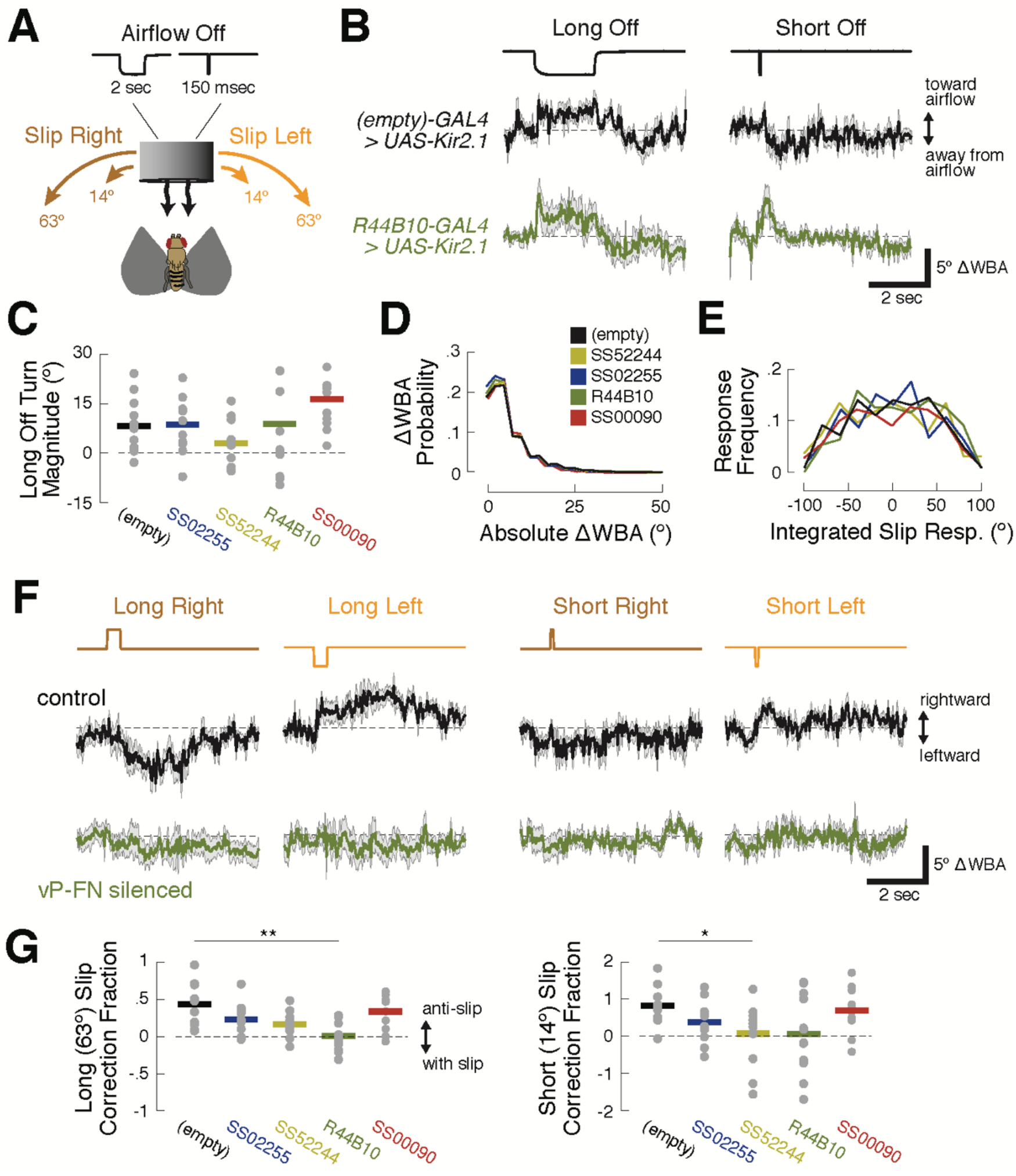
R44B10 neurons are required to convert airflow orientation changes into heading-appropriate turns. (A) Stimulus manipulations. Six manipulations were presented pseudo-randomly every 20 seconds during closed-loop flight: short wind pause (150 msec); long wind pause (2 sec); short (14.44°) and long (63.36°) rightward slip of virtual orientation; and short and long leftward slip of virtual orientation. Slip velocity was 144 °/sec. (B) Responses to long and short airflow pauses in ventral P-FN-silenced flies (*R44B10-GAL4>UAS-Kir2.1*, dark green) and control flies (*empty-GAL4>UAS-Kir2.1*, black). Traces show mean +/- SEM difference in wingbeat angles (ΔWBA), a proxy for intended turning, for 120 trials across 12 files (10 repetitions per fly). In this plot, positive ΔWBA values indicate turns towards the airflow source and negative ΔWBA values indicate turns away from the airflow source. Dashed line represents no turning. (C) Mean ΔWBA (integrated over 2 sec) in response to a long airflow pause for each fly (gray dots) of each genotype. Horizontal bars indicate cross-fly means. Positive ΔWBA values represent turns toward the airflow source. All groups are statistically indistinguishable by rank-sum test. (D) Probability distributions of ΔWBA values for control (black), P-F1 N3-silenced (gold), P-F2N3-silenced (dark blue), all ventral P-FN-silenced (dark green), and E-PG silenced (red) flies. The distributions are statistically indistinguishable by KS-test. (E) Probability distributions of integrated slip responses for each genotype (colors as in (D)). Slip responses are integrated over 5 sec of slip stimulus, with both leftward (negative) and rightward (positive) slips included. The distributions are statistically indistinguishable by KS-test. (F) Responses to orientation slips in ventral P-FN-silenced flies (*R44B10-GAL4>UAS-Kir2.1*, dark green) and control flies (*empty-GAL4>UAS-Kir2.1*, black). Each trace represents the mean +/- SEM of 120 trials across 12 files (10 repetitions per fly). In this plot, positive ΔWBA values indicate rightward turns and negative ΔWBA values indicate leftward turns. Traces show cross-fly mean +/- SEM with colors as in (B). (G) Fraction of slip displacement corrected by each fly (gray dots) of each genotype in response to long (left) and short (right) slips. Values represent mean integrated slip response divided by negative slip magnitude. Positive values indicate “corrective” turns in the opposite direction of the slip. A correction fraction of 1 indicates that a fly steered the airflow direction to be identical before and after a slip trial. Note differing Y-axis scales for each plot. *, *p* < 0.05; **, *p* < 0.01 (rank-sum test).

We next wondered if the poor orienting ability of ventral P-FN-silenced flies arises from motor deficits. However, we found no differences between the distributions of wingbeat angle differences for control and silenced flies (Fig. 7D). Similarly, silencing ventral P-FNs did not change the distribution of integrated turn angles following airflow direction slips (Fig. 7E). Together with our airflow pause data, these results suggest that silencing ventral P-FNs leaves both sensory and motor function intact.

In contrast, silencing ventral P-FNs dramatically impaired the selection of turns following airflow direction slips. In response to slips, control flies generally made corrective turns in the direction opposite the slip (Fig. 7F, top). For rightward slips, control flies made turns to left, and for leftward slips, turns to the right. The duration and magnitude of the slip response varied with slip duration in control flies, such that turns following long slips (Fig. 7F, left) were larger than turns following short slips (Fig. 7F, right). On average, control flies’ reactive turns corrected for 45% of the orientation change induced by long slips, and 90% of short slips (Fig. 7G). Conversely, flies with both types of ventral P-FNs silenced (*R44B10-Gal4>UAS-Kir2.1*) showed no turning response, on average, to slips of any direction or duration (Fig. 7F&G). When only one type of ventral P-FN was silenced, we observed a smaller reduction in mean slip responses. Silencing E-PGs did not disrupt slip correction (Fig. 7F&G), as expected based on the orientation behavior. Together, these results suggest that ventral P-FNs may be specifically involved in generating an appropriate turning response following a change in the direction of airflow.

## Discussion

### Distinct sensory representations in different CX compartments

All animals make use of many different sensory cues to navigate through their environments. Insects and rodents use visual landmarks to return to remembered locations (Ofstad et al., 2011; Collett et al., 2001; Etienne et al., 1990). Odor cues are widely used to navigate towards sources of food or mates (Baker et al., 2018). Movements of the air or water are prominent cues for orientation and navigation across species (Chapman et al., 2011; Alerstam et al., 2015). How neural circuits are organized to process and combine these diverse cues is a fundamental question in neuroscience and evolution.

Although previous studies have examined CX responses to a wide variety of sensory cues (Heinze and Homberg 2007; Weir and Dickinson, 2015; Ritzmann et al., 2008; Phillips-Portillo, 2012; Kathman and Fox, 2019), it has remained unclear to what extent these responses are organized or specialized across CX compartments. By targeting specific cell populations using genetic driver lines, the present study supports the hypothesis that distinct cell types within the CX represent specific sensory cues. Ventral P-FNs exhibited directional responses to airflow that were much stronger than those of the other cell types in our survey. In our study, only E-PGs showed strong directional tuning to the visual landmark, although it is possible that our survey, which only examined responses to 4 cardinal directions, may have missed responses primarily tuned for directions 45° off from these.

A striking finding of our study is that the same sensory cue may be represented in different formats in different CX compartments (Fig. 8). For example, a recent study found a map-like representation of wind direction in E-PGs, with different directions systematically represented across columns (Okubo et al. 2020). This representation is derived from a set of ring (R1) neurons that carry wind information from the LAL — silencing R1 neurons abolishes most wind responses in E-PGs (Okubo et al. 2020). Other sets of ring neurons also carry visual signals to E-PGs (Omoto et al., 2017; Fisher et al. 2019; Turner-Evans et al. 2020). Thus, in the EB, ring neurons appear to carry landmark signals of diverse modalities that collectively anchor the E-PG heading representation, while P-ENs provide angular velocity signals that rotate this representation in the absence of landmark cues (Green et al., 2017; Turner-Evans et al., 2017; Turner-Evans et al. 2020). This model is consistent with our finding that airflow direction cues were not strongly represented in P-ENs, although they are prominent in E-PGs.

**Figure 8.**
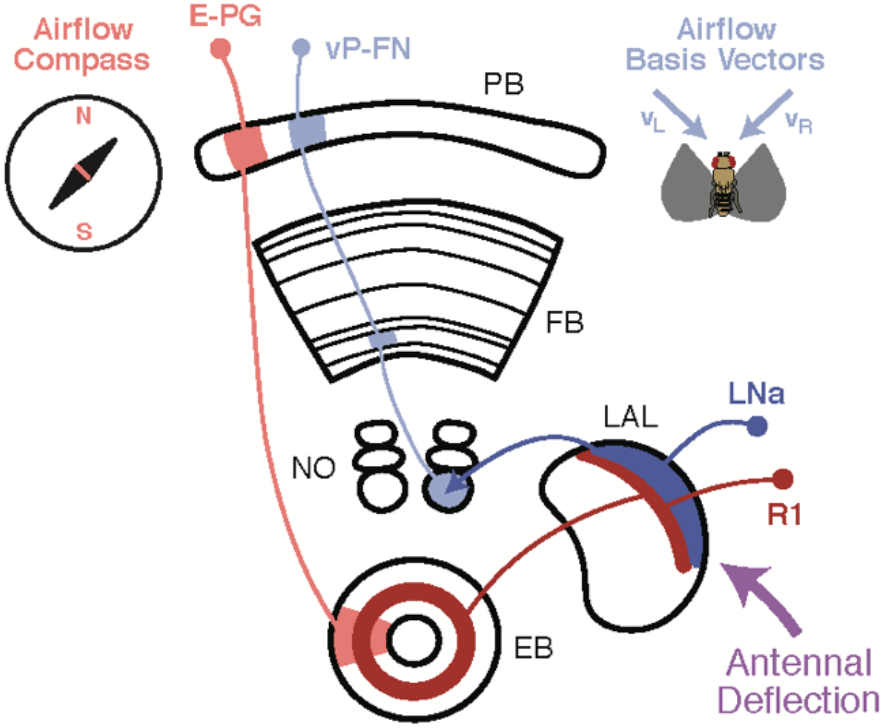
Airflow-representing circuits in the CX. Airflow direction is transduced via antennal deflection signals (purple), which are transmitted through the AMMC and Wedge to the LAL (Yorozu et al. 2009, Suver et al. 2019, Okubo et al. 2020). A recent study (Okubo et al., 2020) found that bilateral antennal deflection signals in the LAL are transmitted to the EB via ring neurons (R1, dark red). R1 neurons are required for formation of an airflow “compass” in E-PG neurons (light red), which represent all possible airflow directions across CX columns. In this study, we showed that LNa neurons (dark blue, 1 per side) are preferentially excited by ipsilateral airflow. LNa neurons carry airflow information from the LAL to the third comparment of the ipsilateral NO (Wolff and Rubin 2018). Ventral P-FNs with cell bodies in one hemisphere each receive input in the third compartment of the contralateral NO. Consistent with this anatomy, ventral P-FNs (light blue) represent airflow from only two directions (appx. 45° to the left and right of midline) across CX columns. Thus, airflow information encoded in the LAL appears to be routed to two different parts of the CX, where it contributes both to a heading compass representation in the EB, and a basis vector representation in the FB.

In contrast to the E-PG airflow “compass”, we found that ventral P-FNs represent airflow as a set of basis-vectors tuned to two orthogonal directions, each originating ~45° to the left or right of the fly midline. Based on our imaging data, we think this representation is most likely inherited from LNa neurons, which similarly represent airflow along two orthogonal axes. This organization of airflow information strongly resembles that of optic flow signals in bee TN1 neurons, which similarly connect the LAL to the NO and provide input to PFN-type neurons (Stone et al. 2017). Thus, the LAL-to-PFN system may represent flow information from various modalities, organized as sets of orthogonal basis vectors. Intriguingly, the compass-like and basis set airflow representations in the EB and FB are likely to arise from a common input pathway in the LAL (Fig. 8). This branching of airflow information may reflect the fly’s need to consider sensory signals in both allocentric (compass) and egocentric (basis vector) reference frames.

### A role for the FB in sensory orientation

Although the CX is broadly required for complex sensory navigation, the precise behavioral role of different CX compartments is still not clear. Martin et al. (2015) found that electrode stimulation at many locations in the FB can produce reliable walking trajectories, suggesting a fairly direct role in locomotor action selection. In contrast, silencing of EB compass neurons (E-PGs) does not impair basic sensory orienting to a visual landmark, but only orienting at a fixed offset (Giraldo et al. 2018, Green et al. 2019). These studies support a model in which different CX compartments, such as the EB and FB, support different aspects of navigation behavior.

Here we found that silencing two classes of ventral P-FNs, but not E-PGs, impaired stable orientation to airflow. When we silenced ventral P-FNs, flies still turned toward the airflow source at flow off, suggesting that these cells are not required to detect airflow or to determine its direction. Flies with silenced ventral P-FNs also exhibited a normal range of motor behavior, arguing that ventral P-FNs do not generate the pool of possible responses to changing airflow direction. Instead, our data suggest that ventral P-FNs guide selection from this pool, specifically converting mechanically detected changes in orientation into an appropriate turning response. These results are broadly consistent with the idea that the FB participates more directly in basic sensory orienting (Honkanen et al. 2019). A caveat is that our strongest effects were observed using a broad line (44B10-GAL4) that labels both classes of ventral P-FNs, as well as some other cells in the CX and other regions. Thus, it remains possible that the more striking phenotype observed in this line arose from off-target expression. However, qualitatively similar effects were observed when only one class of ventral P-FN was silenced. We did attempt to silence several other lines that broadly label ventral P-FNs, but we were unable to find such a line that was viable. Experiments using temporally restricted silencing may help resolve this issue.

### The role of ventral P-FNs in natural behavior

Although we have shown that silencing ventral P-FNs impairs airflow orientation in a tethered flight paradigm, the role they might play in free flight is unclear. During flight, a steady-state wind does not displace mechanoreceptors, but rather displaces the fly (Reynolds et al., 2010; Leitch et al., 2020), generating a strong optic flow signal (Mronz and Lehman, 2008; Theobald et al., 2010) that is often offset from a fly’s heading direction. However, gusts or sudden changes in wind direction can transiently activate antennal mechanoreceptors in flight, leading to behavioral responses (Fuller et al., 2014). Knowing the true direction of the wind in flight would be useful to the fly, both to control dispersal (Leitch et al., 2020), and to orient upwind towards an odor source in flight (van Breugel et al., 2014). Because ventral P-FNs receive input in the protocerebral bridge (PB), presumably carrying heading information from the compass system (Franconville et al., 2018), they may be well-poised to perform this computation. Alternatively, ventral P-FNs might be involved in estimating the direction or speed of self-motion for the purposes of course-control or memory formation (Stone et al., 2017). Future experiments investigating the interaction of airflow, optic flow, and heading signals in these neurons, as well as experiments silencing these neurons during free flight, will provide insight into their function during more natural behaviors.

A final question is how ventral P-FNs are able to control steering to influence orientation to airflow. A small number of descending neurons (DNs) that participate in control of steering during flight have been identified (Schnell et al., 2017; Ferris and Maimon, 2018), although a larger number of DNs target wing motor regions and presumeably play a role in flight control (Namiki et al., 2018). The pathways connecting the CX to these DNs have not yet been elucidated. Ventral P-FNs make most of their outputs in the FB, where they synapse onto a large number of FB local neurons and FB output neurons (Xu et al. 2020). Many of these FB local neurons also receive input from tangential FB inputs, which may carry varied sensory signals. This arrangement could allow sensory inputs such as odor to influence orientation to airflow (Alvarez-Salvado et al., 2018; van Breugel et al., 2014), or for wind cues to be ignored if stronger visual cues are present (Müller and Wehner, 2007; Dacke et al., 2019). Future work aimed at identifying CX output pathways will be critical for understanding how ventral P-FNs and other CX neurons influence ongoing locomotor activity.

## Acknowledgements

We would like to thank Michael Reiser for flies and Michael Long, Gaby Maimon, David Schoppik, and members of the Nagel and Schoppik labs for feedback and helpful discussion. This work was supported by grants from the NIH (R01DC017979 and R01MH109690), and NSF (IOS-1555933) to KIN, a McKnight Scholar Award to KIN, and a New York University Dean’s Fellowship to TAC.

## Author Contributions

Conceptualization – TAC & KIN; Methodology – TAC & KIN; Software – TAC & AMMM; Investigation – TAC & AMMM; Validation – TAC & AMMM; Formal Analysis – TAC & AMMM; Writing – Original Draft – TAC; Writing – Review & Editing – TAC, AMMM & KIN; Supervision – KIN; Funding Acquisition – TAC & KIN.

## Declaration of Interests

The authors declare no competing interests.

## Methods

### CONTACT FOR REAGENT AND RESOURCE SHARING

Further information and requests for resources and reagents should be directed to and will be fulfilled by the Lead Contact, Katherine Nagel.

### EXPERIMENTAL MODEL AND SUBJECT DETAILS

#### Fly Stocks

All flies were raised at 25°C on a cornmeal-agar medium under a 12-hour light/dark cycle. Flies for patch experiments were aged 1–3 days after eclosion before data collection, and flies for behavior experiments were aged 3-5 days. All data shown are from female flies. Parental stocks can be found in the key resources table. Specific genotypes presented in each figure panel are shown below. SS lines contain genetic inserts on chromosomes II and III – the transgenes occupying the second copies of these chromosomes are shown in parenthesis.

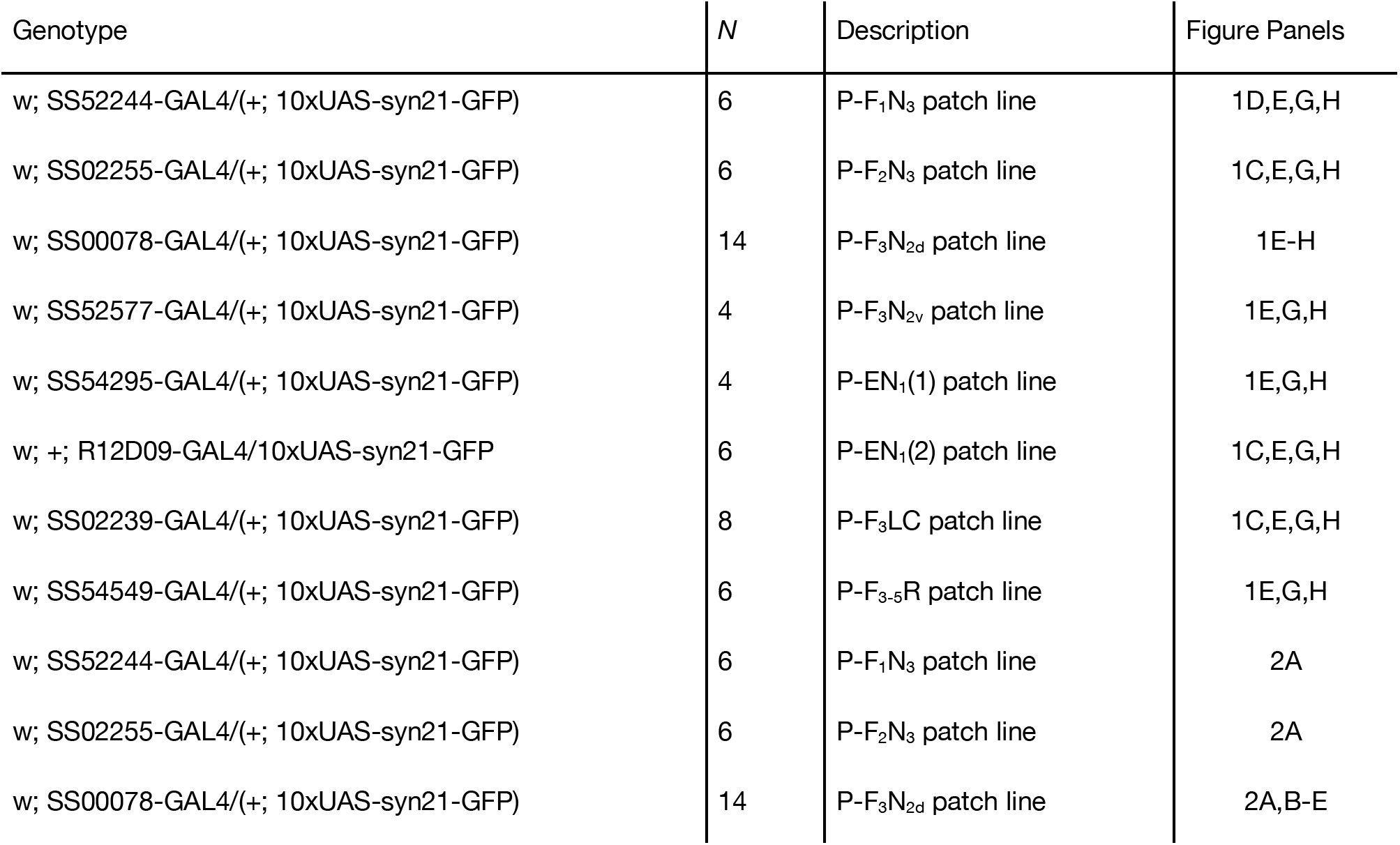
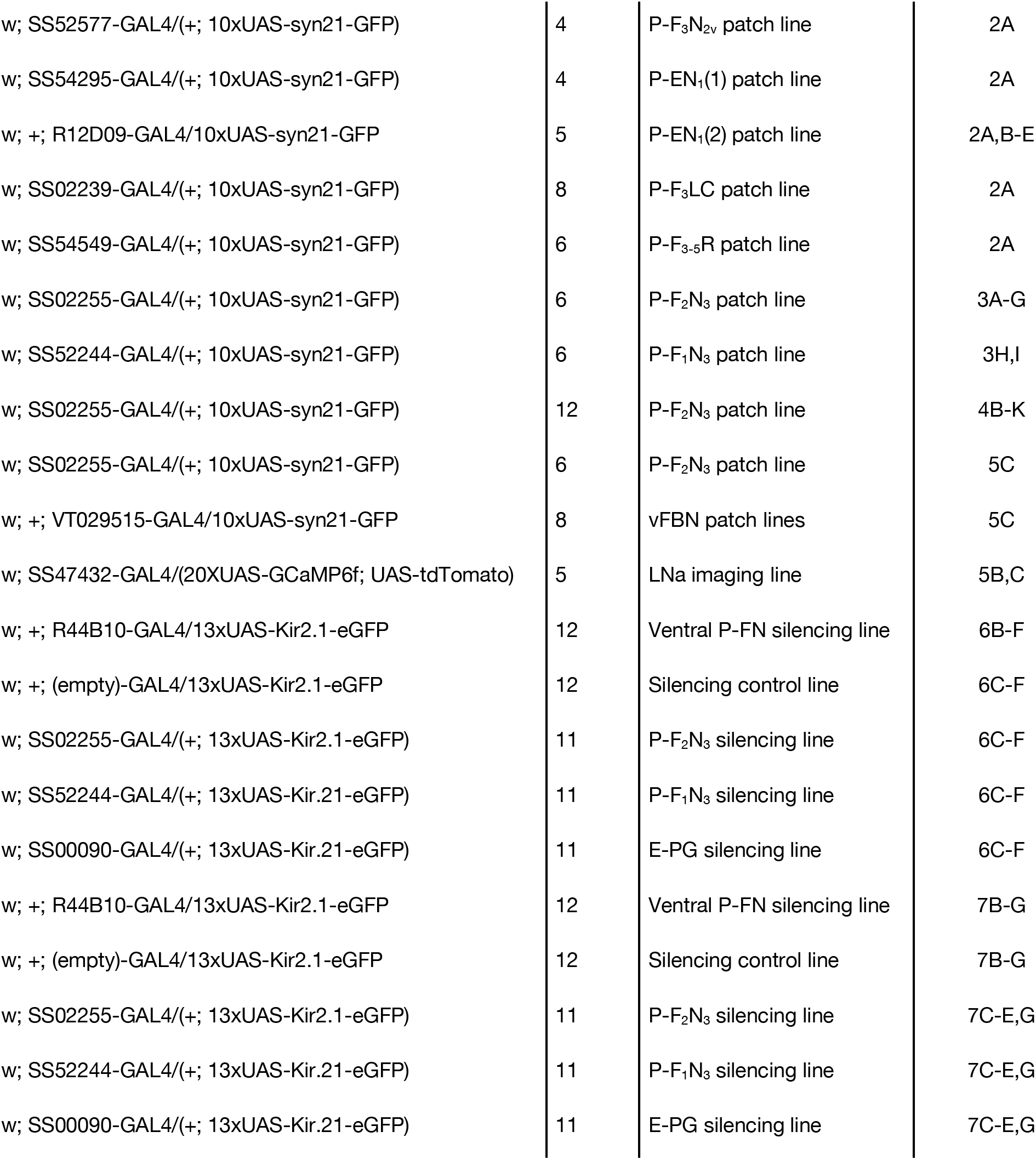

### METHOD DETAILS

#### Electrophysiology

Flies were prepared for electrophysiology (Figs. 1–4) by tethering them to a custom fly holder and reservoir (modified from Weir et al., 2014, see supplemental files for fly holder schematic). All flies were cold anesthetized on ice for approximately 5 minutes during tethering. During anesthesia, we removed the front pair of legs from each fly to prevent disruption of electrophysiology data. We then used UV glue (KOA 30, Kemxert) to fix flies to the holder by the posterior surface of the head and the anterior-dorsal thorax. Once flies were secured, they were allowed to recover from anesthesia for 30-45 minutes in a humidified chamber at 25°C. Prior to patching, we filled the fly holder reservoir with Drosophila saline (Wilson and Laurent, 2005, see below) and dissected away the cuticle on the posterior surface of the head, removing trachea and fat lying over the posterior surface of the brain.

Tethered and dissected flies were then placed in a custom stimulus arena on a floating table (Technical Manufacturing Corporation, 63-541) with a continuous flow of room-temperature *Drosophila* saline over the exposed brain. Briefly, *Drosophila* saline contained 103 mM sodium chloride, 3 mM potassium chloride, 5 mM TES, 8 mM trehalose dihydrate, 10 mM glucose, 26 mM sodium bicarbonate, 1 mM sodium phosphate monohydrate, 1.5 mM calcium chloride dihydrate, and 4 mM magnesium chloride hexahydrate. The solution was adjusted to a pH of 7.2 and an osmolarity of 272 mOsm.

Brains were imaged under 40X magnification (Olympus, LUMPLFLN40XW) by a microsocope (Sutter, WI-SRE3) controlled by a micromanipulator (Sutter, MPC-200). Real-time brain images were captured by a camera (Dage-MTI, IR-1000) and sent to an LCD monitor (Samsung, SMT-1734). Target neurons were identified based on expression of cytoplasmic GFP (see “Fly Stocks” above). Fluorescent stimulation was provided by an LED light source and power controller (Cairn Research, MONOLED). A dichroic/filter cube (Semrock, M341523) allowed for stimulation and emission imaging through the same objective.

We first cleared away the neural sheath overlaying target neurons by puffing 0.5% collagenase-IV (Worthington, 43E14252) through a micropipette (World Precision Instruments, TW150-3) and applying gentle mechanical force. Cell bodies overlaying target somata were then removed via gentle suction, if necessary. Once target neurons were cleaned of debris, we used fire-polished micropipettes (Goodman and Lockery, 2000) to record one neuron per fly. Prior to use, pipettes (World Precision Instruments, 1B150F-3) were pulled (Sutter, Model P-1000 Micropipette Puller) and transferred to a polishing station equipped with an inverted light microscope and a pressurized micro-forge (Scientific Instruments, CPM-2). Pipettes were polished to a tip diameter of 0.5-2 μm and an impedance of 6-12 MΩ, depending on the target cell type. Patch pipettes were filled with potassium-aspartate intracellular solution (Wilson and Laurent, 2005), which contained 140 mM of potassium hydroxide, 140 mM of aspartic acid, 10 mM of HEPES, 1 mM of EGTA, 1 mM of potassium chloride, 4 mM of magnesium adenosine triphosphate, and 0.5 mM of trisodium guanine triphosphate. We also added 13 mM of biocytin hydrazide to the intracellular solution for post-hoc labeling of recorded neurons. The solution was adjusted to a pH of 7.2 and an osmolarity of 265 mOsm. Before use, we filtered this intracellular solution with a syringe-tip filter (0.22 micron pore size, Millipore Millex-GV).

Recorded neurons were confirmed to be of the targeted type in three ways: (1) presence of a fluorescent membrane “bleb” inside the pipette after sealing onto the cell; (2) loss of cytoplasmic GFP through diffusion over the course of the recording session; (3) post-hoc biocytin fill label matching the known anatomical features of the targeted cell type. Two of these three criteria must have been met in order for a patched neuron to be considered a member of a given cell type.

Hardware for electrophysiology was adapted from one previously used (Nagel and Wilson, 2016). In brief, recorded signals passed through a headstage (Axon Instruments, CV 203BU), a pre-amp (Brownlee Precision, Model 410), and an amplifier (Molecular Devices, Axopatch 200B) before being digitized for storage (National Instruments, BNC-2090A) and gain-corrected. Data was collected at 10,000 Hz.

After all electrophysiology experiments, we removed flies from their holders and dissected out their central brains. Dissected tissue was then fixed in 4% paraformaldehyde for 14 minutes at room temperature. Fixed tissue was stored at 4°C for up to 4 weeks before further processing (see “Immunohistochemistry,” below).

#### Stimulus Delivery

Once an active recording was obtained, we presented a series of sensory stimuli from multiple directions. Stimulus delivery was achieved by a modified version of a perviously used system (Currier and Nagel, 2018). Briefly, custom LabView (National Instruments) software controlled the tiggering of airflow (25 cm/s), odor (20% apple cider vinegar), and/or ambient illumination (15 μW/cm^2^) of a high contrast vertical bar that subtended approximately 30° of visual angle. Stimulus intensities for airflow, odor, and light were measured with a hot-wire anemometer (Dantec Dynamics MiniCTA 54T42), a photo-ionization detector (Aurora Scientific, miniPID 200B), and a power meter (ThorLabs, PM 100D and S130C), respectively. All stimuli emanated from the same location in the arena, and the entire arena could be rotated with a stepper motor (Oriental Motor, CVK564FMBK) around the stationary fly. This setup allowed us to present cues from any arbitrary direction. Rotations of the motor were slow (20°/s) and driven at minimal power to minimize vibration and electromagnetic disturbances.

For our initial survey of CX columnar neurons (Figs. 1–3), we used a pseudorandom session design broken down by stimulus direction (−90°, 0°, 90° and 180°) and type (stripe only, airflow only, airflow & stripe together, airflow & odor together, or all 3 stimuli simultaneously). Each 12-second trial included 4 sec of pre-stimulus baseline, 4 sec of stimulus presentation, and 4 sec of post-stimulus time. The first 1 sec of each trial’s baseline period included a 500 ms injection of −2 pA to monitor input resistance over time (this period is not plotted in any Figures). Between trials, a 9-second inter-trial-interval allowed the cell to rest while the motor rotated the arena to the next trial’s stimulus direction. All 20 unique combinations of stimulus direction and condition were presented 4 times each, and each stimulus was presented before the next round of repetitions began. The total session recording time was approximately 50 minutes. If cell health was observed to decay before the session was complete, data was discarded after the preceding “set” of 20 stimuli. For a cell to be included in the survey, at least 40 trials (2 sets of repetitions) must have been completed. Of the 52 neurons patched in the survey, 46 remained healthy for all 80 trials.

To investigate how airflow responses varied by column (Fig. 4), we used 8 directions (−135°, −90°, −45°, 0°, 45°, 90°, 135°, 180°) and presented only the airflow stimulus. Trial and pseudorandom session design were the same as above, but we increased the number of stimulus repetitions to 5, for a total of 40 trials. All 12 flies in this dataset completed all 40 trials.

#### Immunohistochemistry

Fixed brains were processed using standard immunohistochemistry protocols. Briefly, we blocked for 30 minutes at room temperature in phosphate buffered saline (PBS, Sigma, P5493-1L) containing 5% normal goat serum (Vector Laboratories, S-1000) and 0.1 % Triton X-100 (Sigma, X100-100ML). The primary antibody solution was identical to the blocking solution, but had a 1:50 dilution of rabbit anti-GFP antibodies (Fisher Scientific, A-6455) and a 1:50 dilution of mouse anti-bruchpilot antibodies (Developmental Studies Hybridoma Bank, nc82-s). The secondary antibody solution was similarly based on the blocking solution, but also contained a 1:250 dilution of alexa488-conjugated goat antirabbit anitbodies (Fisher Scientific, A-11034), a 1:250 dilution of alex633-conjugated goat anti-mouse antibodies (Fisher Scientific, A-21052), and a 1:1000 dilution of alexa568-conjugated streptavidin (Fisher Scientific, S-11226). Antibody incubations were for 24 hours at room temperature. We washed brains in 0.1 % PBS-Triton three times for 5 minutes after each antibody phase. Immuno-processed brains were mounted on slides (Fisher Scientific, 12-550-143 and 12-452-C) and imaged under a confocal fluorescence microscope (Zeiss, LSM 800) at 20X magnification (Zeiss, W Plan-Apochromat 20x).

#### Calcium Imaging

For calcium imaging, flies (age 10 to 16 days) were anaesthetized and mounted in a simpler version of our electrophysiology holder. Flies were starved for 18-24h prior to beginning the experiment. The back cuticle of the head was dissected away using fine forceps and UV glue was applied to the fly’s proboscis to prevent additional brain movement. Flies were allowed to recover for 5 minutes prior to imaging. The holder chamber was filled with *Drosophila* saline (as above) and perfused for the duration of imaging.

2-photon imaging was performed using a pulsed infrared laser (Mai Tai DeepSea, SpectraPhysics) with a Bergamo II microscope (Thorlabs) using a 20x water-immersion objective (Olympus XLUMPLFLN 20x) and ThorImage 3.0 software. Laser wavelength was set to 920nm and power at the sample ranged from 39-203 mW. Emitted photons were spectrally separated using two bandpass filters (red, tdTOM: 607/70nm, green, GCaMP: 525/50nm) and detected by GaAsP PMTs. The imaging area of approximately 132 × 62 uM was identified using the tdTOM signal. Imaging was performed at 5.0 frames per second.

Airflow and odor stimuli were delivered using a fixed 5-direction manifold (Suver et al. 2019) and controlled by proportional valves (EVP series, EV-05-0905; Clippard Instrument Laboratory, Inc. Cincinnati, OH) using custom Matlab code running on its own PC. We used a hot-wire anemometer (Dantec Dynamics MiniCTA 54T42) to verify that airspeed (~30 cm/s) was equivalent from all 5 directions and did not change during odor delivery. Odorant (apple cider vinegar) was diluted to 1:10 in distilled water on the day of the experiment. Stimuli consisted of 10 s of airflow, 10s of airflow plus odor, followed by another 10 s of airflow with 5 s before and after stimulus presentation. The order of airflow direction was randomized in each block of five trials and we performed 5 blocks per fly. After each block the imaging frame was re-adjusted to account for any drift, and gain and power level were optimized. One fly was excluded from the final analysis as it failed to respond to any stimuli. All flies included include all trials from all 5 blocks.

#### Behavior

For flight simulator experiments (Figs. 5&6) we fixed flies in place using rigid tungsten tethers (see Currier and Nagel, 2018). All flies were cold anesthetized for approximately 5 minutes during the tethering process. During anesthesia, a drop of UV-cured glue was used to tether the notum of anesthetized flies to the end of a tungsten pin (A-M Systems, # 716000). Tethered flies’ heads were therefore free to move. We additionally removed the front pair of legs from each fly to prevent disruption of wing tracking (see below). Flies were then allowed to recover from anesthesia for 30-60 minutes in a humidified chamber at 25°C before behavioral testing.

Tethered behavior flies were placed one at a time in a custom stimulus arena described in a previous paper (Currier and Nagel, 2018). The dark arena was equipped with a pair of tubes (one flow, one suction) that could create a constant stream of airflow over the fly. A camera (Allied Vision, GPF031B) equipped with a zoom lens (Edmund Optics, 59-805) and infrared filter (Midwest Optical, BP805-22.5) was used to capture images of the fly in real-time. Custom LabView software was used to detect the angle of the leading edge of each wing. We multiplied the difference between these wing angles (ΔWBA) by a static, empirically verified gain (0.04, see Currier and Nagel, 2018) to determine each fly’s intended momentary angular velocity. This signal was sent to a stepper motor (Oriental Motor, CVK564FMAK) which rotated the airflow tubes around the fly. The difference in wingbeat angles and integrated heading were also saved for later analysis. This process was repeated at 50 Hz.

Each fly’s 20-minute behavioral testing session was broken down into 20-sec trials that began with a manipulation to the airflow stimulus. Air flowed continuously throughout the entire session, except when interrupted as a stimulus manipulation. We used two durations of airflow pause: 2 samples (100 ms) and 100 samples (2 sec). Additional manipulations included open-loop (not fly-controlled) rotations of the airflow tubes to the left or right of its current position. These open-loop rotations were driven continuously at the maximum speed of the motor (144 °/s) for either 5 samples (14.4°) or 22 samples (63.36°). This gave four additional manipulations: long and short rotations to the left and right. These six stimulus manipulations were pseudorandomly presented ten times each. All flies shown in Figs. 5&6 completed all 60 trials.

### QUANTIFICATION AND STATISTICAL ANALYSIS

#### Analysis of Physiology Data

All data was processed in MATLAB (Mathworks, version 2017B) with custom analysis scripts. Spike times were found by first high-pass filtering the raw membrane potential signal with a second order Butterworth filter (40 Hz cutoff frequency) and then identifying cell type-specific threshold crossings. We used the −2pA test pulse at the beginning of each trial to calculate and track input resistance over time (Fig. S1D).

To calculate membrane potential and spiking responses to our stimuli, we first defined the baseline period for each trial as a 1-second long data segment ending 500 ms before stimulus onset. We additionally defined a response period, which began 500 ms after stimulus onset and similarly lasted 1 sec. Spiking and membrane potential responses in each trial were defined as the mean firing rate or membrane potential during the response period minus the mean of the baseline period. Mean responses (Figs. 1–4 and S2) for each cell to each stimulus were calculated by averaging these responses to stimulus repetitions.

Cross-condition correlation coefficients (Figs. 2, 3 and S4) were found by first taking the mean PSTH across stimulus repeats in each stimulus condition. PSTHs were found by convolving spike trains with a 1 sec Hanning window. We next truncated this full trial mean data to only include the stimulus response and offset periods (seconds 4-12 of each 12-sec trial). Truncated mean response timecourses for the four stimulus directions were then concatenated for each sensory condition. We then found the correlation coefficient between these direction-concatenated mean PSTHs for multi-sensory trials (airflow + stripe) versus airflow only or stripe only.

Response-by column analyses included additional metrics. To calculate the mean orientation tuning of each neuron, we first converted the mean spiking response to each airflow direction into a vector, with an angle corresponding to the airflow direction and a magnitude equal to the mean response to that direction. We then calculated the mean vector across the response vectors corresponding to the 8 stimulus directions that we used. The angle of this mean vector is plotted in Fig. 4E.

Airflow response dynamics were examined using the response to ipsilateral airflow. We computed the cumulative sum to the PSTH during the full 4-sec stimulus period, and divided this timecourse by the integral of the PSTH over the entire 4 seconds (Fig. 4J). We additionally found the time when each cell’s cumulative normalized response reached 0.5, or half its total response (Fig. 4K).

#### Analysis of Calcium Imaging Data

Analysis was performed using the CaImAn Matlab package, Image J, and custom Matlab scripts. We used the CaImAn package (Giovanucci et al. 2019) to implement the NoRMCorre rigid motion correction algorithm (Pnevmatikakis et al. 2017) on the red (tdTOM) time series and applied the same shifts to the green (GCaMP6f) time series. Regions of interest (ROIs) were drawn by hand around the left and right projections between the LAL and NO in ImageJ, using the maximum intensity projection of the tdTOM time series. ROIs were applied to all trials. The positioning of ROIs was adjusted by hand using ImageJ on any trials where significant drift occurred. ImageJ ROIs were imported into Matlab using ReadImageJROI (Muir and Kampa, 2015). We calculated ΔF/F for both tdTOM and GCaMP6f time series by dividing the time series by the average fluorescence of the baseline period (first 5s of the trial). We subtracted the change in red from the change in green to correct for any fluctuations in the Z plane that occurred due to brain movement. The ΔF/F in plots refers to ΔF/F_Green_ -ΔF/F_Red_.

#### Analysis of Behavioral Data

Flight simulator data was collected at 50 Hz as the difference in wingbeat angles (ΔWBA, raw behavior — see Fig. 7) and integrated orientation (cumulative sum of the feedback signal to the arena motor, see above). We parsed the integrated orientation data in three ways. First, we converted the data points into vectors with an angle equal to the fly’s virtual orientation on that sample, and a length of 1. The mean orientation vector was then calculated for each fly. Second, we used the length of a this mean orientation vector to evaluate orientation stability. Third, we found the fraction of samples in the integrated orientation distribution that fell between −45° and 45° (the “toward the airflow” arena quadrant). These measures are all plotted in Fig. 6.

To assess flies’ responses to our airflow manipulations, we analyzed ΔWBA during a 6-sec period surrounding the stimulus manipulation (1 sec pre-stimulus and 5 sec post-stimulus). We found each fly’s mean ΔWBA response across 10 repeats of each slip stimulus. We then found the cross-fly mean and SEM (Fig. 7F).

We additionally integrated ΔWBA over the 5 sec following each slip stimulus to calculate an angular “response” to that slip (Fig. 7E,G,H). Slip response magnitudes were placed into 20° bins and collapsed across slip direction for the purposes of plotting response distributions (Fig. 7E). To calculate the fraction of each slip for which flies corrected, we divided integrated slip response on each trial by the negative magnitude of the stimulus slip, then took the mean correction fraction across all trials of a given slip magnitude (long or short, see Fig. 7G).

To assess flies’ responses to airflow pause, we modified the sign of the raw ΔWBA signal. Because airflow generally drives “with the flow” orienting in rigidly tethered flies (Currier and Nagel, 2018), we wanted a direction-invariant measure of with-/away from airflow turning. Accordingly, we altered the sign of ΔWBA on each sample such that turns toward the airflow source were positive, and turns away from the airflow source were negative (normally, positive ΔWBA indicates rightward turns, and leftward for negative ΔWBA). We then calculated cross-fly mean airflow pause responses (Fig. 7B) as described above for the slip stimuli. For long airflow pauses, we additionally integrated ΔWBA over the 2 sec stimulus period (Fig. 7C).

#### Statistical Analysis

All statistical analyses used non-parametric tests corrected for multiple comparisons (Bonferroni method). For single fly data, paired comparisons were made using the Wilcoxon Sign-Rank test (MATLAB function signrank), and unpaired comparisons using the Mann-Whitney U test (MATLAB function ranksum). Cross-fly (per-genotype) distributions were compared using the Kolmogorov-Smirnov test (MATLAB function kstest2). Significance values for all tests are reported in the Figure Legends.

#### Connectomic Analysis

Data from the fly hemibrain connectome (Xu et al., 2020) was visualized using neuPRINT explorer (*neuprint.janelia.org*). We identified P-F_2_N_3_ as PFNa in this dataset on the basis of NO innervation and nomeclature of Wolff et al., 2015. Similarly, P-F_1_N_3_ was identified as PFNm and PFNp. Based on the connectivity data for an example LNa neuron (1508956088), we determined that P-F_2_N_3_ neurons across CX columns receive significant input from LNa neurons. P-F_2_N_3_ neurons specifically receive input from contralateral LNa neurons in the NO.

### DATA AND SOFTWARE AVAILABILITY

All electrophysiology, behavior, and anatomy data will be made publicly available on Dryad on publication. Code for analysis will be made available on Github on publication.

### KEY RESOURCES TABLE

| REAGENT or RESOURCE | SOURCE | IDENTIFIER |
| --- | --- | --- |
| Antibodies |  |  |
| mouse anti-NC82 | Developmental Studies Hybridoma Bank | RRID: AB_2314866 |
| chicken anti-GFP | Thermo Fisher Scientific | PA1-9533 |
| streptavidin Alexa Fluor 568 | Thermo Fisher Scientific | S-11226 |
| anti-mouse Alexa Fluor 633 | Thermo Fisher Scientific | A-21052 |
| anti-chicken Alexa Fluor 488 | Thermo Fisher Scientific | A-11039 |
| Experimental Models: Organisms/Strains |  |  |
| *SS52244-GAL4* | Bloomington Drosophila Stock Center | RRID: BDSC_86596 |
| *SS02255-GAL4* | Bloomington Drosophila Stock Center | RRID: BDSC_75923 |
| *SS00078-GAL4* | Bloomington Drosophila Stock Center | RRID: BDSC_75854 |
| *SS52577-GAL4* | Bloomington Drosophila Stock Center | RRID: BDSC_86625 |
| *SS54295-GAL4* | Bloomington Drosophila Stock Center | RRID: BDSC_86624 |
| *SS02239-GAL4* | Bloomington Drosophila Stock Center | RRID: BDSC_75926 |
| *SS54549-GAL4* | Bloomington Drosophila Stock Center | RRID: BDSC_86603 |
| *SS47432-GAL4* | Bloomington Drosophila Stock Center | RRID: BDSC_86716 |
| *R12D09-GAL4* | Bloomington Drosophila Stock Center | RRID: BDSC_48503 |
| *R44B10-GAL4* | Bloomington Drosophila Stock Center | RRID: BDSC_50202 |
| *(empty)-GAL4* | Bloomington Drosophila Stock Center | RRID: BDSC_68384 |
| *10xUAS-IVS-syn21-GFP-p10 (attP2)* | Michael Dickinson | N/A |
| *13xUAS-Kir2.1-eGFP/TM3* | Michael Reiser | N/A |
| *20xUAS-GCaMP6f* | Bloomington Drosophila Stock Center | RRID: BDSC_42747 |
| *UAS-tdTomato* | Bloomington Drosophila Stock Center | RRID: BDSC_36328 |

## Supplementary Material

**Figure S1.**
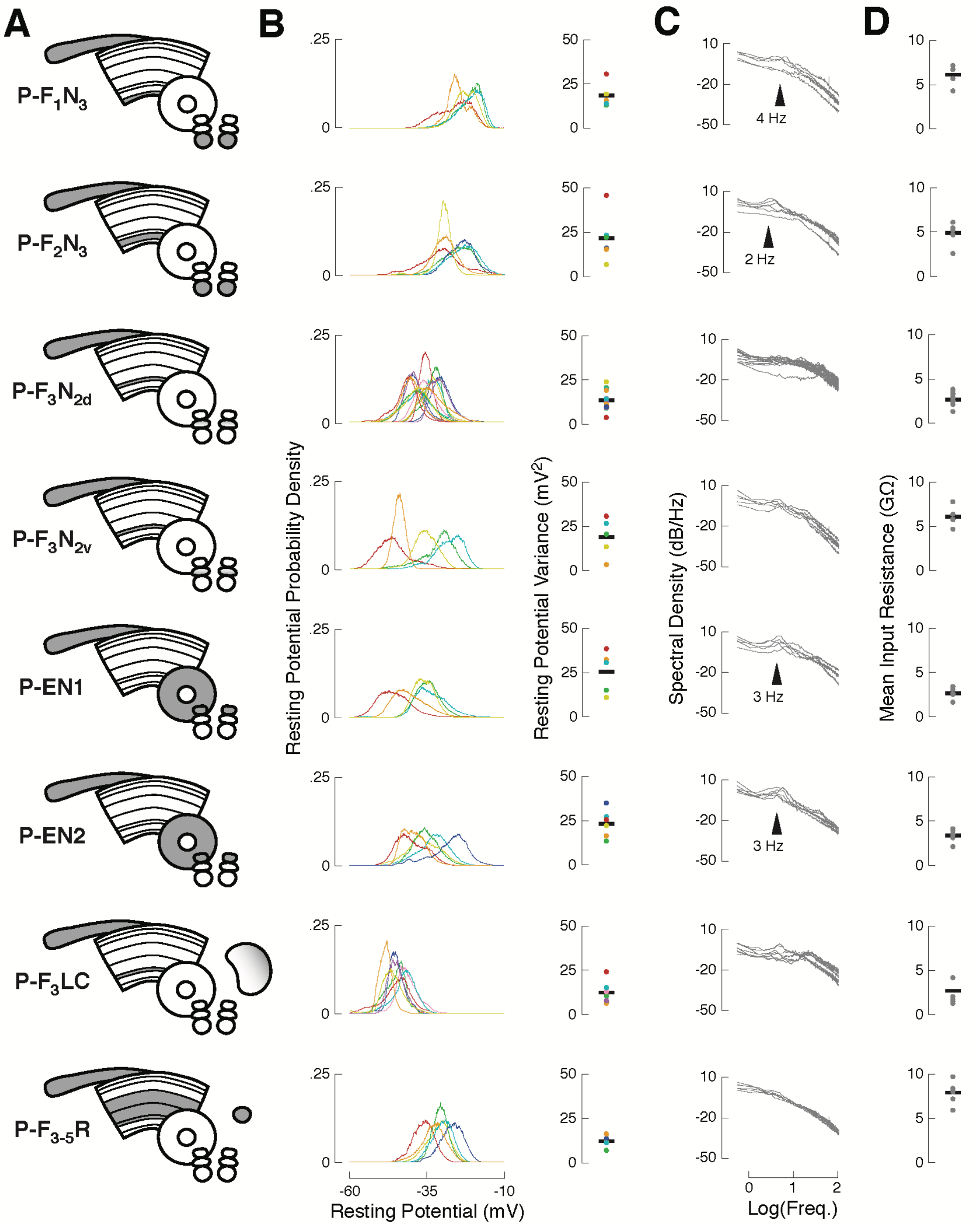
related to Figs. 1–3. Baseline activity characterization for recorded cell types. (A) CX neuropils innervated by each recorded cell type. Each cell type is named after standard nomenclature: single letters represent innervated neuropil (gray), with putative input regions before the dash, and putative output or mixed process regions after the dash. Each recorded neuron was filled with biocytin to confirm its identity. Previous work identified two anatomically identical but functionally distinct classes of P-ENs. PB, Protocerebral Bridge; FB, Fan-shaped Body; EB, Ellipsoid Body; NO, noduli; ROB, round body; LAL, Lateral Accessory Lobe. Each row in panels (B)-(D) contains data from the cell type shown. (B) Left: resting membrane potential distributions for each neuron (colored lines) from the indicated cell type. Ventral P-FNs (P-F1N3 and P-F2N3) rested high (−25 mV), P-F3LC rested low (−50 mV), and all other cell types rested near −35 mV. Right: resting membrane potential variance for each neuron (colored dots) from each cell type. Black bars indicate cross-cell means. P-ENs and some P-FNs showed the greatest potential variance at rest, due to rhythmic/bursty baseline activity (see C). (C) Spectral density of the raw voltage signal for each neuron (gray lines) from the indicated cell type. Note that the x-axis is plotted on a log scale. Noteworthy deviations from 1/F density are marked with arrowheads. Both types of P-EN showed strong rhythmic/burst activity with an interval of approximately 350 msec. Ventral P-FNs displayed less regular rhythmicity at different timescales. (D) Mean input resistance for each neuron (gray dots) from each cell type. Input resistance was sampled on each trial, so each point is the mean of 40-80 Rin measurements, depending on the number of trials completed for each neuron. Black bars indicate cross-cell means.

**Figure S2.**
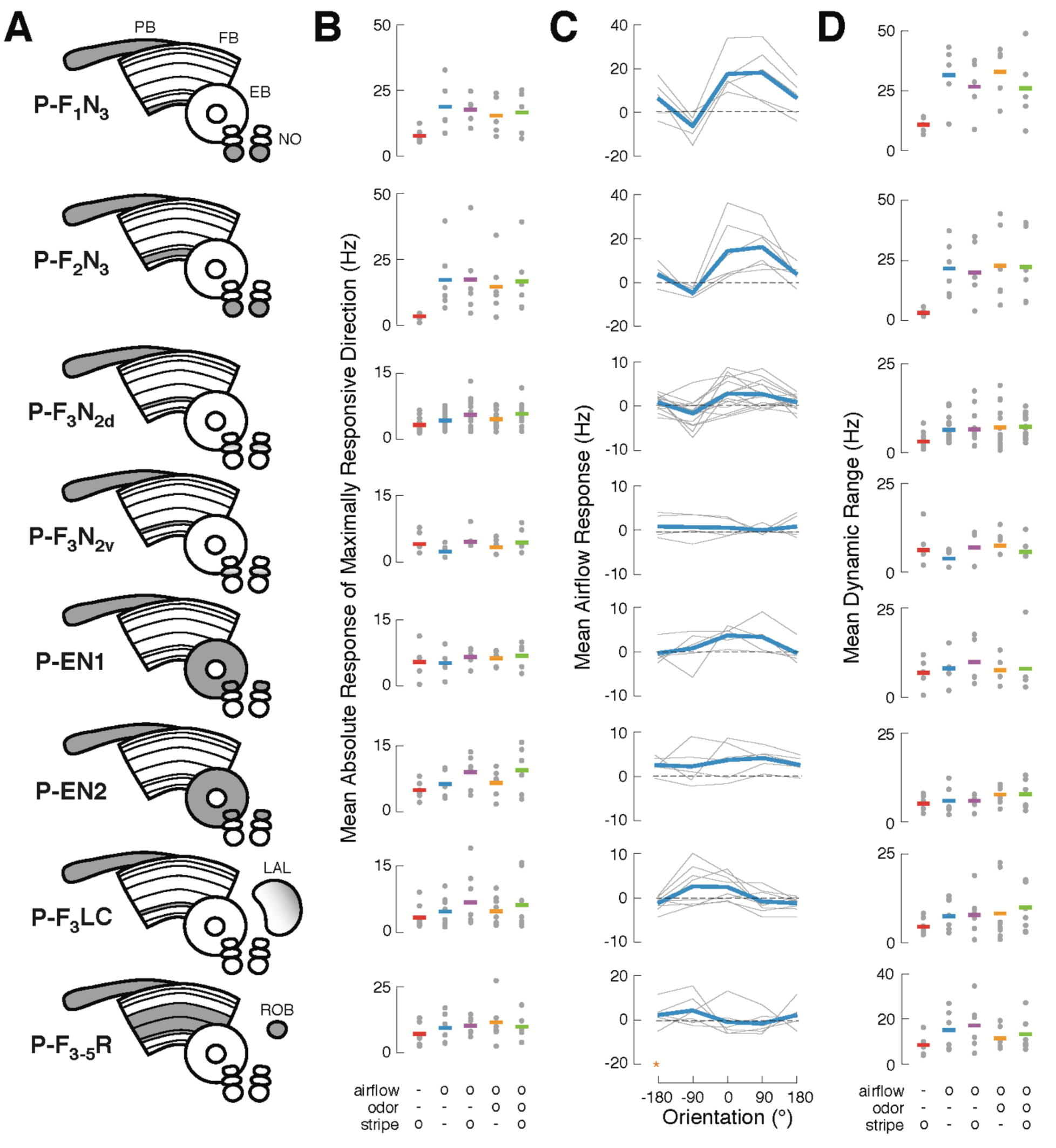
related to Fig. 1. Summary of sensory responses across CX cell types. (A) CX neuropils innervated by each recorded cell type. (B) Maximal responses to each stimulus condition. Gray dots represent the mean absolute spiking response of each cell to four presentations from the direction that elicited the largest response. Horizontal bars represent the mean across cells (colors as in Fig. 1B). All responses represent changes from baseline. Note different vertical scales. (C) Tuning for airflow direction. Mean spiking response as a function of airflow direction for each neuron (gray), and across neurons of a given type (blue). Note different vertical scales. Data at −180° is replotted from 180° for clarity (orange star). (D) Dynamic range for each modality/condition, equal to the difference between the most excitatory and most inhibitory responses across directions. Larger values indicate stronger tuning. Single cells and cross-cell means as in (B).

**Figure S3.**
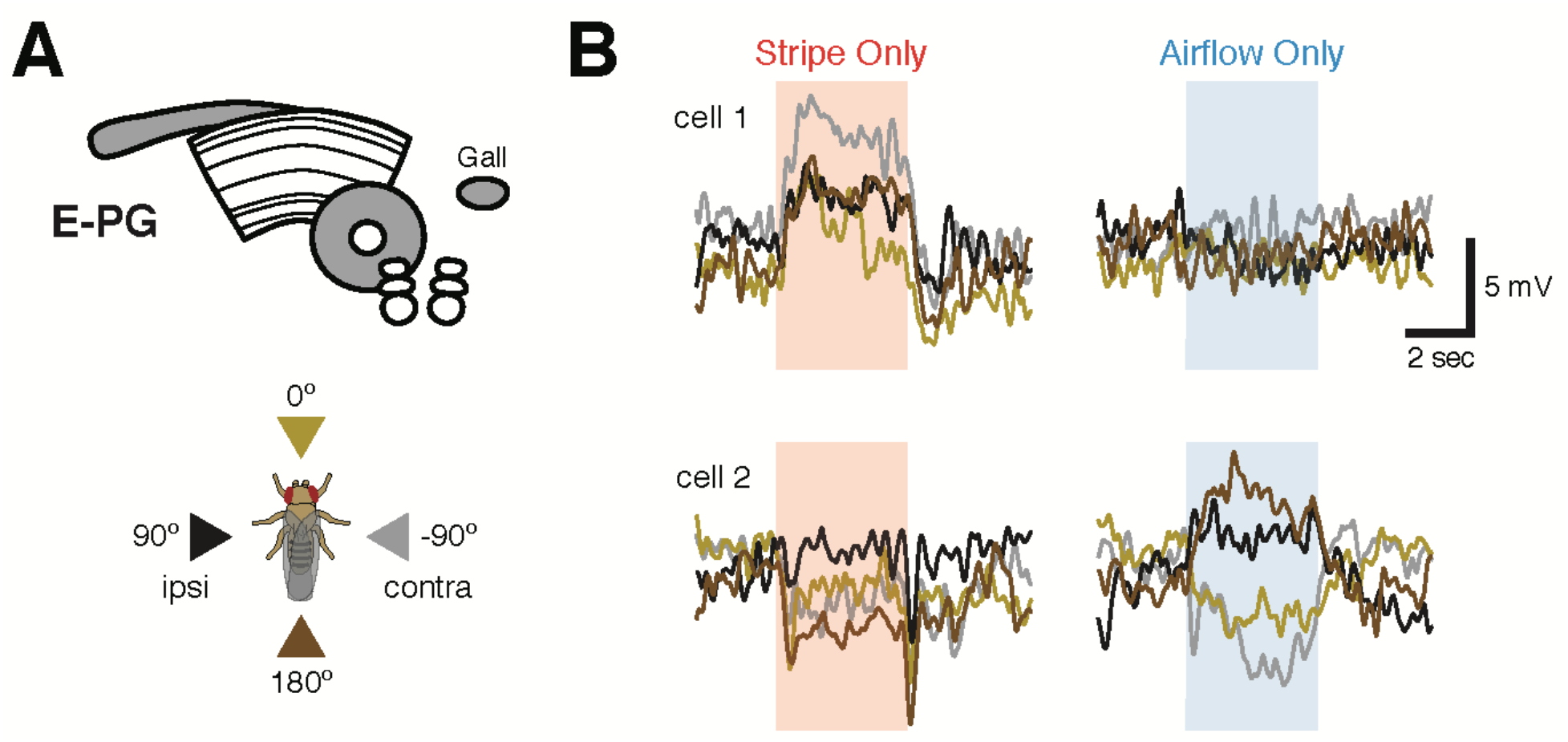
related to Fig. 1. Tuned visual responses in E-PGs. (A) Top: neuropil schematic of E-PG (“compass”) neurons, which are known to be tuned for both visual landmark orientation (Green et al., 2017) and airflow direction (Okubo et al., 2020). Bottom: stimulus direction color key. (B) Membrane potential responses to stripe or airflow for two example E-PG neurons. Each trace is the mean of four presentations of a stimulus from a single direction. Traces are colored according to the direction schematic shown in (A). Single E-PG neurons shown directional tuning for stripe, airflow, both, or neither. Neurons that appear untuned for a stimulus likely possess off-cardinal preferred directions not captured in our stimulus set. Note that preferred airflow and stripe directions may not be identical for a single E-PG.

**Figure S4.**
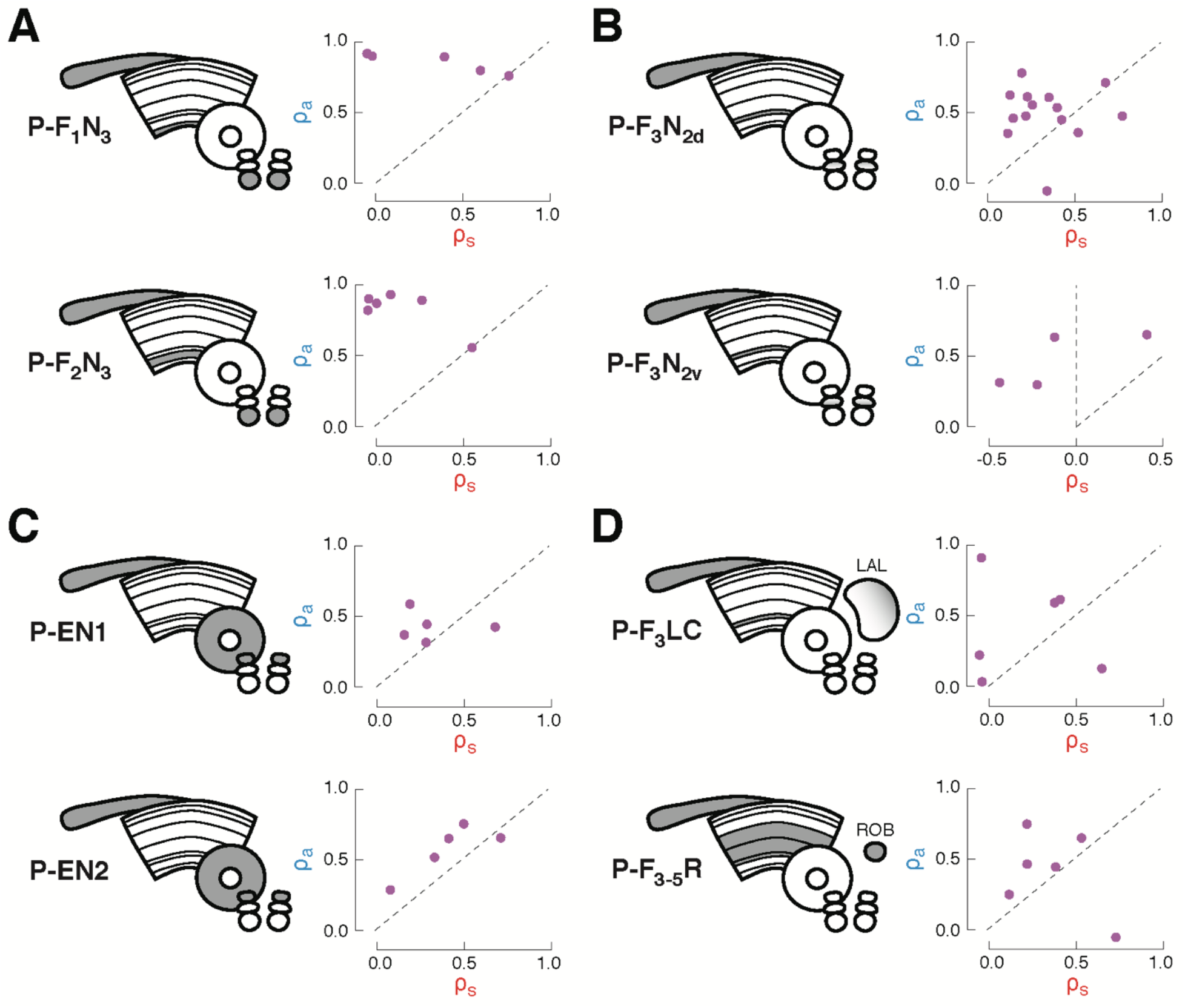
related to Figs. 2&3. Characterization of multi-sensory integration for recorded cell types. Each plot show the similarity (correlation coefficient) of the response to airflow + stripe with the response to airflow alone (y-axis) versus stripe alone (x-axis), as in Figs. 2&3. Each point represents the coefficient for one fly, calculated from the mean firing rate timecourses for all four stimulus directions. Data along the diagonal indicates that the multi-sensory response is equally similar to the stripe alone and airflow alone responses, a hallmark of summation. Data above the diagonal indicates that the multisensory response more closely resembles the airflow response and data below the diagonal indicates that the multisensory response more closely resembles the stripe response. (A) Ventral P-FNs, P-F1N3 and P-F2N3, consistently possess high airflow coefficients, indicating that the multi-sensory response is dominated by the airflow response. (B) Dorsal P-FNs, P-F3N2d and P-F3N2v, show diverse coefficients, indicating diverse integration strategies across cells. P-F3N2v neurons even exhibit strong negative coefficients (left of vertical dashed line), indicating that the multi-sensory response can resemble inverted single modality responses. (C) P-ENs. P-EN1 integrates variably across cells, while P-EN2 integration is more consistent, as discussed in Fig. 2. (D) CX extrinsic columnar neurons. Integration across P-F3LC and P-F3-5R neurons was the most variable among recorded cell types.

**Figure S5.**
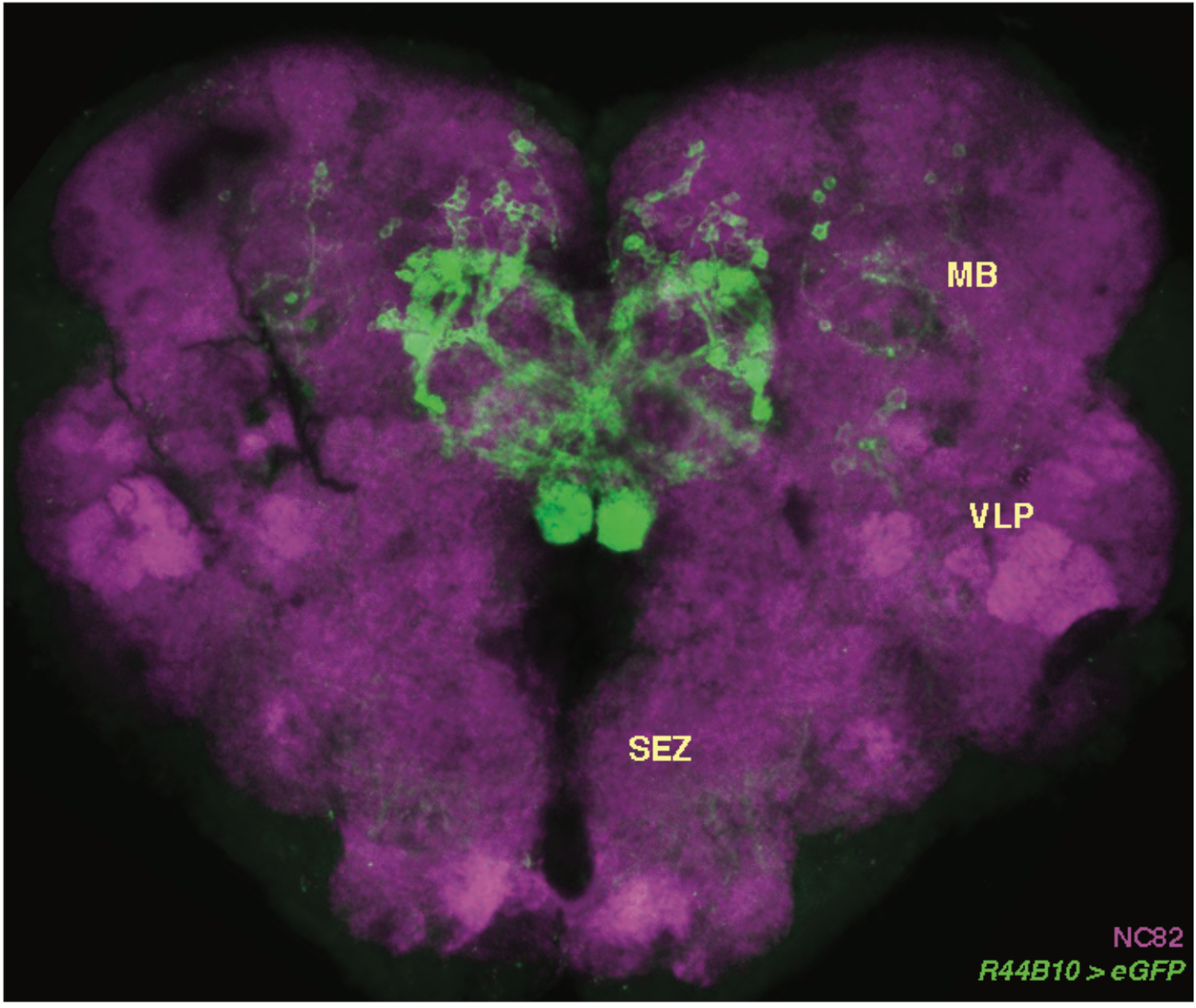
related to Figs. 5&6. Full central brain anatomy of *R44B10-GAL4*. Little GFP expression can be seen outside of the CX in *R44B10-GAL4*. Weak off-target signal is present in the mushroom bodies (MB), ventrolateral protocerebrum (VLP), and sub-esophageal zone (SEZ). These regions are not labeled in the split-GAL4 lines we used. Neuropil in magenta.

## Notes

### Competing Interest Statement

The authors have declared no competing interest.

